# Anthropogenic land use exerts selection pressures on the resistome of a wild rodent

**DOI:** 10.64898/2025.12.12.693898

**Authors:** M. Gicquel, A. Planillo, E. Heitlinger, S.K. Forslund-Startceva, S. Kramer-Schadt, S. C. M. Ferreira, V. H. Jarquín-Díaz

**Affiliations:** Department of Ecological Dynamics, Leibniz Institute for Zoo and Wildlife Research (IZW), Alfred-Kowalke-Straße 17, 10315, Berlin, Germany; Institute for Biology. Department of Molecular Parasitology. Humboldt University Berlin (HU). Philippstr. 13, Haus 14, 10115, Berlin, Germany; Federal State Agency for Consumer & Health Protection Rhineland-Palatinate, Koblenz, Germany; Max-Delbrück-Center for Molecular Medicine in the Helmholtz Association (MDC), Berlin, Germany; Charité – Universitätsmedizin Berlin, corporate member of Freie Universität Berlin and Humboldt-Universität zu Berlin, Experimental and Clinical Research Center, Lindenberger Weg 80, 13125 Berlin, Germany; Experimental and Clinical Research Center, a cooperation between the Max-Delbrück-Center for Molecular Medicine in the Helmholtz Association and the Charité - Universitätsmedizin Berlin, Germany; Institute of Ecology, Technische Universität Berlin, Rothenburgstr. 12, 12165 Berlin, Germany; University of Veterinary Medicine Vienna, Research Institute of Wildlife Ecology, Veterinärplatz 1 1210, Vienna, Austria

## Abstract

Antimicrobial resistance poses a significant global health challenge. However, the factors maintaining antimicrobial resistance genes (ARGs) in wildlife microbiomes remain unclear particularly in species inhabiting human-dominated landscapes. We analysed 875 gut metagenomes from wild house mice (*Mus musculus*) on German farms between 2016 and 2022 to identify environmental and host determinants of ARG occurrence. Using joint species distribution models, we quantified the influence of landscape, climate and mouse associated characteristics on the occurrence of individual ARGs and on trait dependence among genes. Environmental variables and livestock farming intensity explained more than 21% of ARG variation, whereas host characteristics accounted for less than 10%. Analysis of ARG traits revealed that agricultural land use and exposure to livestock increased the occurrence of mobile ARGs. Pig density was strongly associated with integron-encoded sulfonamide resistance genes and genes conferring tetracycline and beta-lactam resistance. Consistent with these findings, mouse resistomes closely resembled those of livestock manure. Taken together, our results show that landscape conditions, particularly farming intensity, shape the distribution of specific ARGs and mobile ARGs in house mice microbiomes. Understanding the factors impacting ARGs prevalence in wildlife is crucial for determining transmission of antimicrobial resistant microorganisms from animal and environmental reservoirs.

## Background

Antimicrobial resistance (AMR) represents a major concern for global health responsible for an estimated 5 million deaths worldwide yearly^1,2^. In response to this crisis, the World Health organisation and the Food and Agriculture organisation established the “Global Action Plan on Antimicrobial Resistance” in 2015, aiming for a coordinated One Health approach including human, animal and environmental health sectors^3^. The complexity of transmission routes, origins and reservoirs within and between humans, farm animals, and their shared environments poses challenges for implementing effective control strategies^4^. In particular, our understanding of how different environmental factors impact the occurrence or potential transmission of antibiotic resistance genes (ARGs) in bacterial communities associated with wildlife hosts or the external environment, which both act as AMR reservoirs, is still limited^5,6^.

The mechanisms of antimicrobial resistance are a natural phenomenon that have enabled bacteria to survive, persist and evolve within specific ecological contexts, which are determined by the dynamics of the microbial community^7,8^. The use of antimicrobial drugs in medical, agricultural and livestock production imposes additional evolutionary selection pressure on bacterial communities, driving the occurrence and transmission of ARGs^9–13^. Human activities have accelerated the horizontal spread of established ARGs to clinically relevant pathogens even from evolutionarily distant bacteria through mobile genetic elements^14^. Latent genes represent, in counterpart, a large collection of less studied genes conferring resistance that remain understudied at environmental bacterial communities and largely absent from databases^15^. Under antibiotic selection pressure, ARGs can also evolve from precursor genes, so called *proto-ARGs*, that have other functions within bacterial genomes and do not directly confer a resistance phenotype^16–18^. Thus, the resistome of a bacterial community is constituted by the metagenomic composition of established, latent and precursor genes, all of which may directly or indirectly contribute to resistance. The resistome varies substantially between environments, which represents important implications for the emergence and dissemination of resistance.

Metagenomic screening has extensively characterised the resistome of human gut, wastewater and livestock^19–22^. Such high throughput approaches have been instrumental to identifying socio-economic, genetic and environmental factors associated with resistance potential, and the dynamics of clinically relevant ARGs from environmental reservoirs to humans^10,23–26^. Although some wildlife species are recognised reservoirs of infectious diseases and antimicrobial resistance, systematic metagenomic screening of wildlife associated resistomes is still sporadic. This limits our understanding of background levels of resistance in species able to mobilise antimicrobial resistant bacteria over large geographical distances or to maintain local dissemination^27,28^. Therefore, the use of high-throughput approaches in wildlife species becomes crucial in two directions: first, to determine the basal resistome in their microbiomes and second to assess the impact of environmental drivers in resistance dynamics.

In human-dominated landscapes, generalist wildlife species able to move freely at the interface between natural and anthropogenic environments encounter diverse sources of ARGs, e.g. via foraging, and may act as “mobile links” for resistance dissemination^29–31^. While birds and large mammals have been highlighted as sentinels for resistance due to their capacity for long-distance movement, small mammals, particularly rodents, are key to understanding ARG transmission at local scales, and for distinguishing environmental effects in transmission and spread^32–34^. Rodents, such as rats (*Rattus norvegicus*, *R. rattus*) and house mice (*Mus musculus*), living near human settlements and domestic animals, as well as wild rodents that have never been exposed to antibiotics, are all known to carry ARGs and resistant bacteria in their gut^35–37^ . The latter highlights both the role of “pest” species as carriers and disseminators of resistant bacteria and ARGs, and the widespread nature of antibiotic resistance in the ecosystem. In the context of antimicrobial resistance ecology, it is crucial to understand how bacteria carrying ARGs interact with their “total environment”, i.e. their direct host “environment” as well as the wider landscape context. Microbial communities and their functional potential including resistance genes are shaped by different host characteristics, e.g. different life-history strategies such as dispersal or philopatry or sex differences^38^. In this respect, it is also of importance how ecological dynamics within rodents’ gut microbiota are impacted by external environmental changes and could be used to detect, predict and even control the emergence and spread of resistance threats.

Changes in the environment, land use and landscape structure are expected to influence wildlife and their associated microbial communities through different biotic and abiotic factors that surround them. Especially farms with high livestock density or extensive crops are sources of bacteria carrying ARGs and pollute other environments through manure, dust moved by wind, water or even wildlife^39^. Spatially-explicit integration of environmental factors along environmental gradients in a multidimensional way provides a framework to understand how microbial communities and their resistomes respond to environmental change and which factors contribute to the composition of ARG traits like mobility^40^. Nevertheless, the linkage from environment via host to microbiome to resistome has been rarely incorporated to investigate how abiotic environmental factors in an ecosystem shape the distribution and diversity of ARGs in “wild” microbial communities due to imperfect integration of ecological frameworks into microbiome research.

Here, we analysed the resistome of densely sampled wild house mice (*Mus musculus*) communities across transects spanning hundreds of kilometers in Germany. Under a spatially-explicit approach, we assessed whether the composition, potential mobility and selection of ARGs within gut microbial communities are associated with anthropogenic, environmental and climatic factors, and disentangled these effects from host-specific influences. Our study provides an integrative landscape scale perspective on the transmission routes and environmental drivers underlying ARG dynamics in mobile wildlife hosts.

## Material and Methods

### Study area

We conducted our study in Germany, central Europe, in the federal states of Berlin, Brandenburg, Bavaria and Mecklenburg-Vorpommern (Figure 1). Human population density in Germany is around 241 people/km^2^, with 77% of the population living in urban areas (www.worldometers.info). The states selected for the study area are characterised by a large proportion of land dedicated to agriculture, from circa 70% in Mecklenburg-Vorpommern to 45% in Brandenburg and Bavaria, and 4% in Berlin (https://www.laiv-mv.de/Statistik/, https://www.statistikdaten.bayern.de/, Amt Für Statistik Berlin-Brandenburg, 2025). Forested areas occupy 24%, 37%, 36% and 18% of the state surface, respectively. Within agriculture, a large proportion is dedicated to animal husbandry, with Germany being the main European producers of milk or pork (www.bmleh.de). Crops and animal husbandry occur across the country, with different levels for each state. Poultry farming is present in all states, but it is predominant in Mecklenburg-Vorpommern. Cattle farms are found mainly in Bavaria and some areas in Brandenburg. Pig farming occurs in some locations in Bavaria and Mecklenburg-Vorpommern. In contrast, Berlin and Brandenburg produce mainly vegetables (www.bmleh.de).

**Figure 1.**
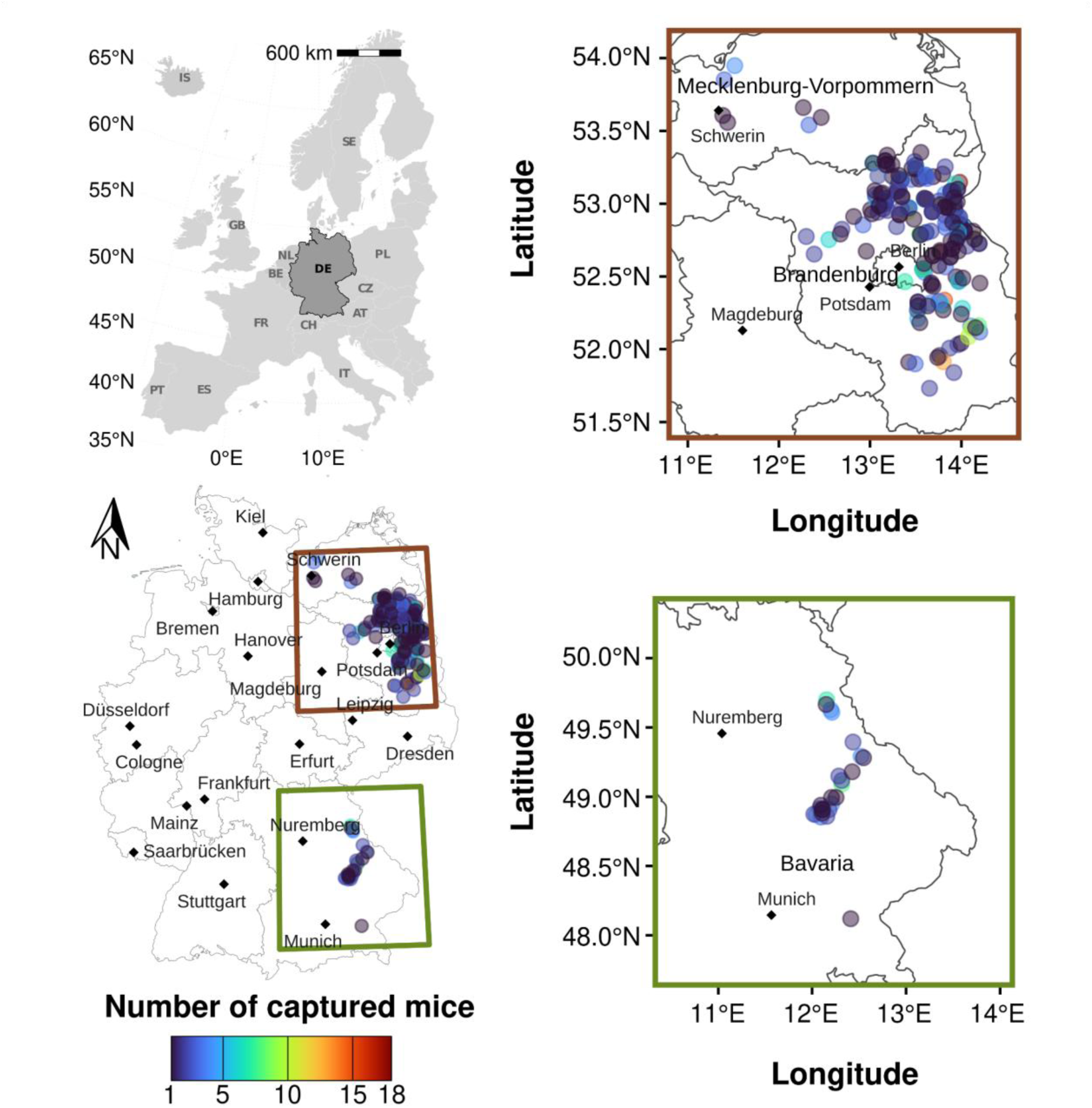
Geographical distribution of farms where mice were captured. The points colour indicates the number of mice captured at each farm location.

### Sample collection

House mice (*Mus musculus;* hereafter: mouse) were captured alive from different private properties (farms) between 2015 to 2022, with only 2020 trapping missing due to COVID-19 contact restrictions. After capture, mice were housed individually in cages overnight and euthanized by overdoses of isoflurane. Animal capture and handling was conducted under the animal experimentation licenses No. 2347/35/2014 and 2347/28/2021 issued by LAVG Brandenburg. All mice were dissected within 24 h after capture. The gastrointestinal colon content was collected, flash frozen in liquid nitrogen and later stored at − 80 °C for DNA extraction at a later time^41^.

### DNA extraction and sequencing

Metagenomic DNA from colon content was extracted in two randomized batches. Samples collected between 2015 and 2018 (N= 679) using NucleoSpin®Soil kit (Macherey-Nagel GmbH & Co. KG, Düren, Germany) following the manufacturer’s protocol with modification as reported in previous studies^38,41^. Samples from 2019, 2021 and 2022 (N= 196) were extracted using the ZymoBIOMICS-96 MagBead DNA Kit (Zymo Research Europe GmbH, Freiburg, Germany) in the automatized system TECAN Fluent® 780 NAP workstation following the manufacturer’s instructions with minimum minimum modifications to the lysis step. In brief, 750 μL ZymoBIOMICS Lysis Solution was added to ∼100 mg of colon content and transferred to ZR Bashing Bead Lysis Tubes with beads (0.1 and 0.5 mm). The samples were mechanically disrupted using a PeQLab Precellys 24 (Bertin Corp., Rockville, MD, USA) for 2×15 s at 5500 rpm. After 5 min rest, the cycle was repeated two more times for a total of three cycles. The DNA was eluted with 50 μL of ZymoBIOMICS DNase/RNase Free Water. Aliquots with at least 750 ng quantified using Qubit fluorometric quantification dsDNA Broad Range Kit (Thermo Fisher Scientific, Walham, USA) were shipped on dry ice for library preparation and sequencing at the Competence Centre for Genomic Analysis (Kiel, Germany). Sequencing libraries were prepared with the Illumina DNA prep kit following manufacturer’s instructions. Shotgun sequencing was done on the Illumina NovaSeq6000 S4 platform (Illumina, San Diego, California, USA) using 300 cycles and v1.5 reagent kits.

### Bacterial taxonomy, ARG annotation, diversity and composition

Paired-end shotgun sequencing reads were assessed for quality using FastQC v0.12.1 and processed using the NGLess pipeline v1.5^42^. Briefly, quality control consisted of trimming and filtering the reads with a minimum quality of 25, a minimum length of 45 b and removing the first 6 nucleotides. Two sequential decontamination steps were then performed. The first was to remove human contamination by filtering reads with a minimum match size of 45 and 90% identity to the reference human genome assembly GRCh38.p14. The second decontamination step aimed to remove host DNA by filtering reads mapped to the reference house mouse (*Mus musculus*) genome assembly GRCm39 - mm39 using the same criteria as for the human genome. Both reference genomes were masked for low complexity regions and regions mapping to the proGenomes3 gene catalogue^43^. The taxonomic profiling of decontaminated metagenomic reads was performed with mOTUs3^44^. Samples with less than 1000 reads assigned to bacteria were removed from the analysis and only mOTUs with abundance above 0.005% and 1% prevalence within each metagenome were included in the analysis. Antimicrobial resistance gene (ARG) annotation to summarise ARG occurrence and abundance in each sample was performed using Resistance Gene Identifier (RGI bwt) v5.2.1-2. Metagenomic reads were aligned to the Comprehensive Antibiotic Resistance Database (CARD) v3.1.4^45^ using the bowtie2 aligner v2.4.5^46^ and annotated following Antibiotic Resistance Ontology (ARO). For purposes of this study, only ARO IDs with more than 80% coverage to the reference were retained for further analysis and denoted as ARGs. ARG abundance was normalised to Fragments Per Kilobase of gene per Million mapped reads (FPKM) to control for differences in sequencing depth between samples and gene length differences. In addition, abundances were binned to ARG, AMR gene family and drug classes level. ARGs were manually regrouped based on the drugs to which they confer resistance. ARGs referring to penam, cephalosporin, carbapenem, cephamycin, penem and monobactam were grouped into the beta-lactam class. Those referring to macrolides, lincosamides and streptogramins were grouped into the MLS class. Those referring to more than one drug class were grouped into the multidrug class. The mobility of the ARGs was determined based on their genetic locations with MGEs by referencing CARD Data. The richness of ARGs observed for each sample was calculated as the total ARG detected in the dataset for a given sample, using Microbiome v1.26.0^47^. Principal component analysis (PCA) and permutational multivariate analysis of variance (PERMANOVA) were performed based on centred-log-ratio (CLR) transformed FPKM ARG abundances and 9999 permutations using ‘adonis2’ from package ‘vegan’ v2.6-475. PERMANOVA for both microbial and ARG profiles was run by = “‘margin” using year of collection, mouse sex, longitude, latitude and transect as geographical references, taxonomically assigned read counts and level of ARG count as predictors, stratifying by sequencing batch. Comparisons of ARG richness were conducted using a two-sided Mann–Whitney U-test with Bonferroni correction for multiple comparisons. The effect size *r* was calculated as the Z statistic of the MWU test divided by the square root of the sample size as implemented in the package rstatix v0.7.2. Significance was determined at a p-value cut-off of 0.05 unless otherwise stated. All figures were created using the following packages: ggplot2 v3.5.1, ggsci v3.2.0, ggpubr v0.6.0, gridExtra v2.3 and RColorBrewer v1.1-3 with minor editions using Inkscape v0.92.5. Data handling was performed using pipelines compatible with tidyverse v2.0.0.

The bioinformatic analysis was performed on the High Performance Computing Cluster from the Max Delbrück Centrum, Berlin (Max-Cluster) equipped with 5.2k Cores – 46TB RAM.

### Retrieval of publicly available livestock manure metagenomes

To assess the proximity of house mice derived resistomes to those from farm animals, we used two available datasets from livestock manure. One from the project EFFORT including pooled manure from pig, cattle and poultry herds in nine European countries^25,48^. The second from rural and urban pig and poultry farms in Ghana, which was included as a non-european reference^49^. A total of 639 fecal metagenomes were downloaded from the NCBI Sequence Read Archive (BioProjects PRJEB22062 and PRJEB62878). Both datasets were generated using Illumina platforms which ensure compatibility with data generated for our study. To avoid any bias in the comparisons, all metagenomes were processed and ARGs annotated in the same way as house mice derived metagenomes (*see Bacterial taxonomy and ARG annotation section*).

### ARG traits and explanatory variables

We annotated detected ARGs based on four traits: i) Mobility (binary) indicates whether the ARG is associated with mobile genetic elements (MGE) as reported in the CARD database (FALSE: Chromosomally encoded ARGs; TRUE: ARG associated with mobile elements such as plasmids, integrons, transposons). ii) Localization (categorical) indicates the type of mobile genetic element in which the gene could be located. In our dataset, localization includes chromosome, plasmid, integron, transposon, pathogenicity island, multiple mobile genetic elements, and mixed when it could be either on the bacterial chromosome or in a MGE. Due to low its representation, multiple MGEs and pathogenicity islands were grouped together. iii) Drug class group (categorical) groups ARGs into broader drug class categories to increase statistical power due to the sparse representation of some individual classes. Grouping was based on similarity in chemical structure (Supplementary Table S1). iv) Resistance mechanism (categorical) is a functional mechanism by which the ARG confers resistance and is classified into antibiotic target alteration, antibiotic inactivation, efflux, target protection, target replacement, and reduced permeability. A subset of ARGs were analysed as clinically relevant based on the World Health Organization’s list of medically important antimicrobials^50^(Supplementary Table S2).

We prepared a set of host-level and environmental explanatory variables hypothesised to influence the distribution of antibiotic resistance genes (ARGs) in house mice. As mouse host characteristics reflecting different life-histories, we used sex (female/male), the body condition *(BC)* of the mice calculated as the residuals of body weight and body length relationship and a mouse density proxy, calculated as the number of trapped individuals per site (*Ntrapped*). While mouse density may influence microbial exchange or exposure risk, the body condition might reflect disturbed microbiomes either as cause or consequence of bad health.

Environmental variables included proportions of different land cover features such as proportion of forest or agriculture in a 100m raster, accessibility of the farm calculated by distances to linear transport features, farm animal density describing municipality-level livestock densities for cattle, pigs, and poultry as well as climatic data, averaged for the month of August in the respective sampling year.

All spatial variables were projected to a common coordinate reference system (ETRS89 / LAEA Europe, EPSG:3035, see source and original specifications of spatial layers in Supplementary Table S3), rasterised if needed and resampled to a 100 × 100 m resolution for spatial consistency. The proportion of agricultural areas was derived from the Corine Land Cover spatial layer class 2 by calculating the percentage of the land use class within each 100 × 100 m grid cell. Distances to roads and paths were calculated by rasterizing the vector map to a 100 m resolution and calculating the distance to each linear road / path feature to the center grid cell coordinates. Values from these layers were then extracted for each sample based on their geographical location. Farm animal information was obtained from the Thünen agricultural atlas as number of livestock units per municipality^51^. We selected the three main livestock species in our study area (cattle, pigs and poultry) and transformed the number of heads of each species into densities to obtain a measure comparable across all sampling sites. The densities were obtained by dividing the number of heads by the total area of the corresponding municipality (Supplementary Figure S4).

To reduce multicollinearity among explanatory variables, we calculated pairwise Pearson correlation coefficients *r* and excluded one variable from each pair with |r| > 0.7, retaining the most ecologically informative predictors (Supplementary Figure S5).

## Data analysis

### Factors affecting ARG distribution and abundance

We applied two Joint Species Distribution Models (jSDM) within the Bayesian framework of Hierarchical Models of Species Communities (HMSC)^52^ to model the occurrence (presence-absence) and the abundance of ARGs as a function of selected explanatory variables. Models were implemented using the *Hmsc* v 3.3-7^53^. The HMSC framework is ideal for the analysis of multiresponse data, as it allows for the inclusion of both fixed and random effects, as well as controlling for the spatial autocorrelation structure in the residuals, while analysing the data at the level of the taxa or the traits. Additionally, it also provides a measure of association between the different taxa after controlling for the effects of all explanatory variables, i.e., positive or negative associations after accounting for environmental preferences.

The response matrix consisted of presence-absence or abundance data, respectively, for ARGs detected in individual mice sampled across multiple farms, municipalities, and regions. To assess how ARG trait–host-environment interactions shaped ARGs distributions, we also included a trait matrix comprising relevant ARG characteristics (see *ARG traits and explanatory variables*) in both models. To model binary ARG presence-absence data, we used a probit error distribution, and to model ARG abundance we log-transformed it (log(x+1)) to fit a Gaussian distribution. For the abundance models, we first filtered out ARGs with an overall prevalence lower than or equal to 5% to avoid chain convergence problems.

After the removal of correlated explanatory variables, we additively included the mouse characteristics (sex, body condition, mouse density), landcover (agricultural land, impervious surface, tree cover density, small woody features), accessibility (distance to read, distance to path) farm animal density (cattle, pig and poultry) and climatic characteristics (precipitation and temperature) as explanatory variables into the model. To account for the nested and spatio-temporal structure of the study design, we specified three levels of random effects: Spatial autocorrelation was accounted for by including the XY-coordinates of farm locations as a spatially-explicit random effect. To account for local farm-level variability and nested effects of the sampling design with multiple mice per farm, we included *FarmID*. Lastly, we included *Year* as a temporal random effect to capture inter-annual variability.

For the ARG occurrence model, we ran four parallel Markov Chain Monte Carlo (MCMC) chains, with 1000 burn-in iterations, followed by 10,000 sampling iterations and a thinning interval of 10, resulting in 4000 posterior samples per parameter. For the ARG abundance model, we ran four parallel Markov Chain Monte Carlo (MCMC) chains, with 1000 burn-in iterations, followed by 15,000 sampling iterations and a thinning interval of 10, resulting in 6000 posterior samples per parameter.

Model convergence was assessed using MCMC trace plots, Gelman-Rubin diagnostics, and effective sample sizes of a random subset of parameters. Model performance was evaluated using Tjur’s R², or R², and the area under the receiver operating characteristic curve (AUC). To quantify the contributions of the host and environmental explanatory variables versus the nested and spatio-temporal structure of the data, we decomposed R² into variance components associated with fixed effects and each random effect using the partitioning tools in the *Hmsc* v 3.3-7.

### Association of house mouse and livestock manure resistomes

After preprocessing and ARG annotation, the resistomes of livestock manure and house mouse were merged into a single dataset for further analysis. The resulting dataset includes FPKM abundances for each ARG, along with country and host of origin and ARG ontology. The combined dataset was compiled using Phyloseq v1.48.0^54^. Resistome richness and dissimilarity were estimated as described for the house mouse dataset alone (*see Bacterial taxonomy and ARG annotation section*). The relative effect of the host of origin on differences in richness against the house mouse resistome was estimated by fitting a linear model with ARG richness as response and host and country as predictor variables using ‘stats’ v3.6.2. Estimated marginal means (EMMs) for each host, and pairwise comparisons between EMMs of mouse and livestock resistome richness were calculated using ‘emmeans’ v1.11.1. Fisher’s exact test was used to determine the degree of intersection in ARGs detected between the resistomes of mice and livestock. Pairwise comparisons were performed between each livestock host and the mouse resistome, and the p-values were adjusted using Bonferroni correction for multiple comparisons. An Euler diagram to represent ARG overlaps between the resistome of different hosts was done with ‘eulerr’ v7.0.2. To determine whether the resistome composition of mice was more similar to that of a particular livestock host, a linear mixed-effects model was fitted using ‘lme4’ v1.1-37. The model used *1 - Aitchison* distance as measure of resistome similarity between pairs of samples. Only resistome similarities within mice and between mice and livestock hosts were used. Host pair and country pair were used as fixed effects, and the sample IDs of the pairs were included as random effects to control for pseudoreplication. As described for the richness model above, EMMs were calculated for each host pair, and the relative degree of resistome similarity was determined by calculating pairwise comparisons between EMMs within the mouse and between mouse-livestock resistomes. The prevalence of each ARG in mice and livestock was estimated as the proportion of metagenomes from each host where the given gene was detected. Comparisons of host-based prevalence were used to determine the potential sources of genes between the livestock hosts and the mice. Genes were categorised as co-promoted if they were present in ≥10% of samples from both mice and a given livestock host. Genes present only in ≥10% of samples from one type of host were categorised as host-specific. ARGs present in fewer than 10% of samples from mice and a given livestock host were classified as non-promoted^55^.

All analyses were performed in R version 4.4.1 (R Core Team, 2024).

## Results

### Composition of the house mouse resistome and microbiome

We detected 340 ARGs in the metagenomes of 875 house mice. The majority of the detected genes (178) potentially confers resistance to the ten relevant antibiotic classes that are of high clinical concern. However, these genes accounted for only 9% of the total abundance. The remaining genes corresponded to non-specific multidrug genes, including mutations in housekeeping genes (e.g. rRNA) and promoter regions, and members of different multidrug efflux pump families, accounting for up to 90% of the total gene abundance (Figure 2a). The most prevalent resistance mechanism is antibiotic target alteration (147 genes), which included 79 multidrug genes and 14 genes associated with other xenobiotics, such as disinfectants and antiseptics. Among the genes conferring resistance to relevant drug classes, those related to the target protection resistance mechanism for tetracyclines, fluoroquinolones, and MLS (macrolides, lincosamides, and streptogramins) antibiotics accounted for 70% of the abundance and were distributed across all years. Only 16% of the genes (55 genes) had prevalence higher than 50% and abundance above 2.5e6 FPKM across mouse metagenomes, including five conferring resistance to tetracycline and one to beta-lactam antibiotics (Supplementary Figure S6). About 23.5% (80 genes) were potentially associated with mobile genetic elements (MGEs). Most of the genes (260) were located on the main bacterial chromosome and correspond to multidrug or mutation in housekeeping genes associated with *Escherichia coli* variants. On average, each mouse metagenome contained 59.6 genes (Figure 2b), with few mice showing ARG richness higher than 100, which remained consistent over the years. During the sampling period of this study, we observed a significant increase in the number of genes detected towards the end; particularly, those from the last two years showed a strong increase. Those from 2016 generally had lower richness (Wilcoxon *p*-value adjusted <0.01, effect size *r* >0.2, Table 1). ARG composition was highly similar between mice; year explained only 2% of the variance and the composition remained consistent over time (PERMANOVA, p-value > 0.05; Figure 2c).

**Figure 2.**
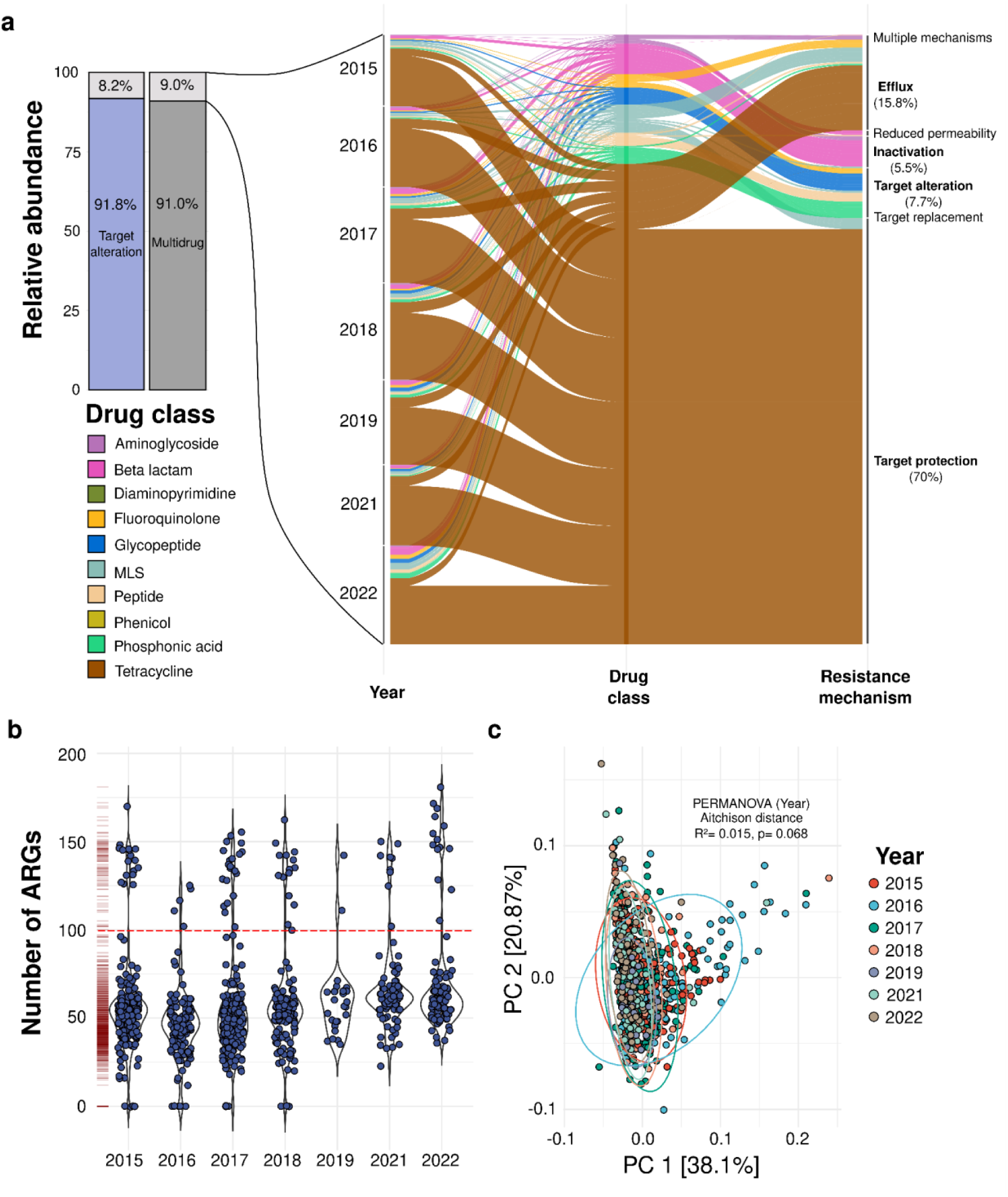
General overview of ARGs in house mouse microbiomes. **(a)** Average relative abundance of ARGs in all house mouse microbiomes from this study. More than 90% of the abundance was related to multidrug genes (139 genes) and target alteration genes (147 genes) as resistance mechanisms. Genes conferring resistance to the ten relevant drug classes account for only 9% of the total abundance. Of these, the majority were genes conferring resistance to beta-lactam antibiotics (46 genes), accounting for less than 1% of the total abundance. Genes conferring resistance to tetracycline (18 genes) represent 6% of the total abundance. Genes conferring resistance to tetracycline, fluoroquinolone and MLS (macrolides, lincosamides and streptogramins) antibiotics, which are related to the target protection mechanism, accounted for 70% of the abundance related exclusively to the ten major antibiotic classes. Genes with efflux, target alteration or antibiotic inactivation each represented more than 5% of the abundance related exclusively to the ten major antibiotic classes. **(b)** ARG richness in house mice over the years. The number of ARGs detected in 2015 and 2016 was significantly lower than at the end of the study period (Wilcoxon, *p*-value adjusted <0.001). Most mice (n = 774) had an ARG richness of around 50 genes, while a smaller group (n = 85) had a higher number of genes detected. The dashed red line defines a threshold of 100 ARGs detected. Each point represents a single mouse metagenome. **(c)** PCA showing dissimilarity in ARG profiles between years of sampling. Each point represents the ARG profile of a sample. Distances between points reflect biological gene composition dissimilarity based on Aitchison distances. Points and ellipses are coloured by year of sampling.

**Table 1.** Differences in ARG richness in house mice at different years.

| Year 1 | Year 2 | N 1 | N 2 | P-value adj. | Significance | Effect size |
| --- | --- | --- | --- | --- | --- | --- |
| 2015 | 2016 | 191 | 127 | 0.0002436 | *** | 0.25 |
| 2015 | 2021 | 191 | 84 | 9.56E-05 | **** | 0.28 |
| 2015 | 2022 | 191 | 104 | 2.84E-05 | **** | 0.28 |
| 2016 | 2018 | 127 | 126 | 0.005061 | ** | 0.23 |
| 2016 | 2019 | 127 | 26 | 0.011865 | * | 0.28 |
| 2016 | 2021 | 127 | 84 | 3.70E-12 | **** | 0.51 |
| 2016 | 2022 | 127 | 104 | 8.55E-15 | **** | 0.54 |
| 2017 | 2021 | 217 | 84 | 7.25E-08 | **** | 0.34 |
| 2017 | 2022 | 217 | 104 | 4.10E-10 | **** | 0.37 |
| 2018 | 2021 | 126 | 84 | 0.0003318 | *** | 0.30 |
| 2018 | 2022 | 126 | 104 | 0.00013713 | *** | 0.30 |

Only significant comparisons are shown

A total of 606 bacterial mOTUs belonging to 50 different families were detected in the house mouse microbiomes. Each metagenomic sample contained an average of 212 mOTUs (range: 25–374), and the composition remained homogeneous over the years, sexes, geographical locations, and levels of ARGs detected in the samples (PERMANOVA, p > 0.05). Members of the *Lachnospiraceae* and *Muribaculaceae* families were highly prevalent (prevalence > 98%) and represented the majority of abundance, at 25.8% [95% CI: 24.7–26.8] and 10.7% [95% CI: 9.99–11.5], respectively. Other bacterial families that could potentially mediate ARG transmission, such as *Enterobacteriaceae* and *Enterococcaceae*, had a prevalence below 50%, or were not detected, e.g. *Staphylococcaceae*. However, *Enterobacteriaceae*, and particularly *Escherichia coli* abundance showed a strong correlation with the ARG richness within each mouse resistome (Spearman’s correlation, Rho > 0.5, *p*< 0.001) and partially explained the difference in ARG levels in some resistomes (Supplementary Figure S7).

### Impact of environmental characteristics on the occurrence of ARG and compositional structure within house mouse microbiomes

To determine the impact of environmental characteristics on the occurrence (presence-absence) of ARGs in house mice microbiomes, we employed joint Species Distribution Models (jSDM) for each of the ARG response, controlling for geographical and temporal proximity between samples and the farm identity and considering selected mouse-related characteristics, environmental factors. The jSDM on ARG presence-absence had an average effective sample size of over 800, which is indicative of robust parameter estimation and low autocorrelation. Additionally, Gelman-Rubin Diagnostic parameters for beta and gamma were higher than 1, indicating adequate model convergence. Overall, the model showed strong predictive performance (AUC= 0.91), identifying samples with high and low probability of specific ARG occurrence (Supplementary Figure S8). On average, 65% of the variance in ARG occurrence in the mouse gut bacterial microbiome was explained by random effects, and 35% by explanatory variables (Figure 3a). The identity of each farm (Random: Farm ID) was the dominant source of explained variance, which indicates strong farm-specific effects not captured by environmental or host factors. Unmeasured ecological and farm management variables at the local level likely influenced the large variance explained by farm. Spatial proximity between samples led to similar ARG occurrence profiles, indicating local spread or spatially similar conditions not included in our models.

**Figure 3.**
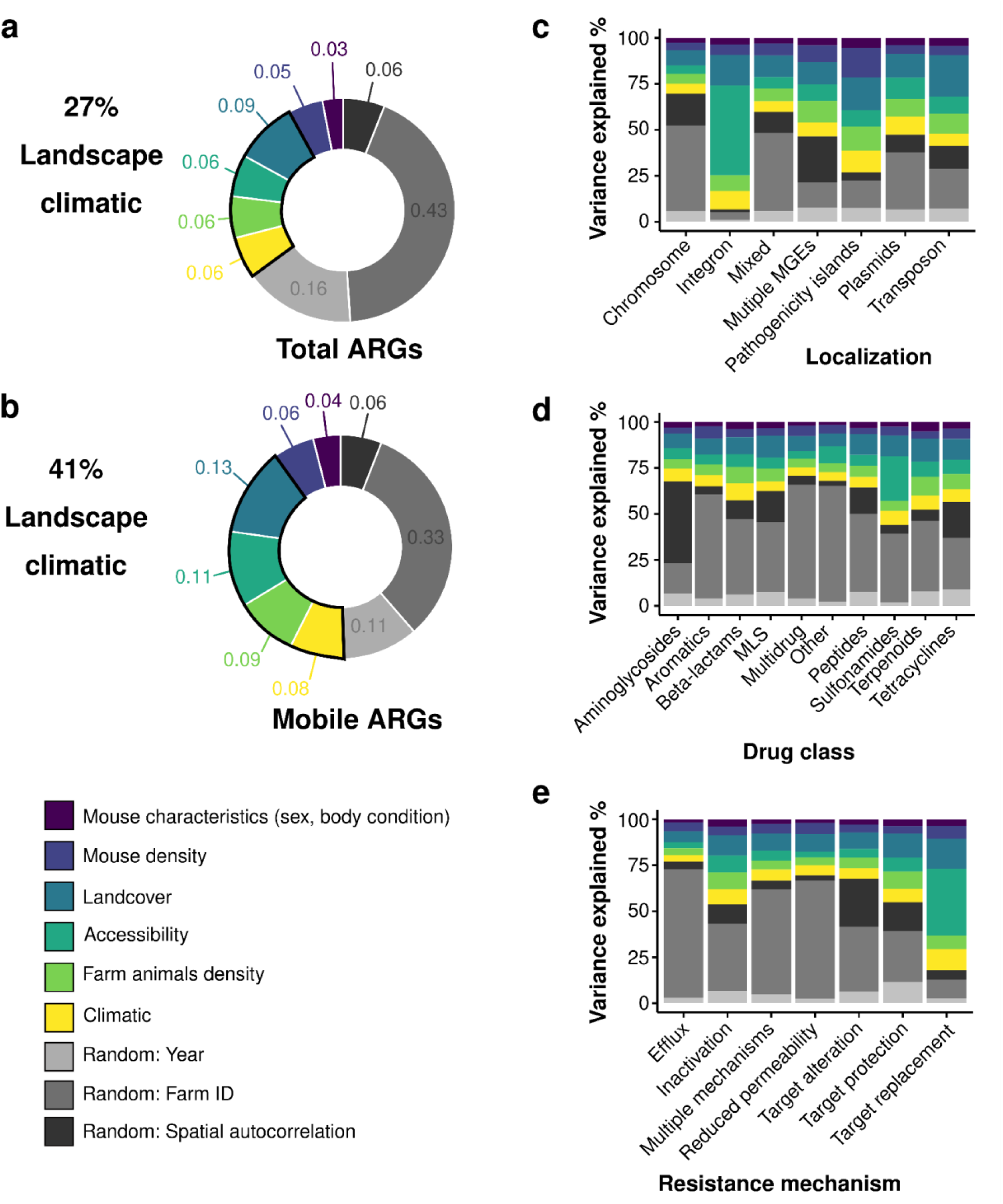
The relative contribution of mouse characteristics, environmental factors and random effects to the variation in the occurrence of ARGs in the mouse gut microbiome. **(a)** Variance partitioning revealed that the farm of origin and the geographical and temporal proximity between the samples (random effects) had a larger impact on the occurrence of all the detected ARGs, accounting for over 60% of the total variation. Environmental factors, including landscape characteristics and climatic factors, accounted for an additional 27%. Mouse characteristics and density explained less than 10% **(b)** When these effects were evaluated only for potentially mobile genes, the environmental factors increased their explanatory contribution to 41% of the ARG occurrence, indicating that mobile genes were less dependent on specific farm characteristics. **(c-e)** The variance partitioning averaged for each ARG’s traits shows the occurrence of genes with potential localization in integrons, genes conferring resistance to sulfonamides and with target replacement mechanisms are more influenced by environmental factors.

Mouse-related characteristics had a low impact in the occurrence of ARG. Mouse density, represented by the number of mice trapped at the farm site, explained 5%, while mouse condition and sex explained only 3%. Among the environmental parameters, land cover represented by agricultural lands, tree cover, small woody features and impervious surfaces explained about 9%. Farm accessibility, together with farm animal density (poultry, pig, cattle) and climatic variables explained each additional 6%.

We used potential localization as a proxy of ARG mobility. Thus, when the model predicted the occurrence of only ARGs in any mobile genetic element, mouse characteristics and environmental factors explained more (51%) than the model for all genes combined (35%). Land cover, accessibility, farm-related variables and climate particularly had greater explanatory power for mobile genes, showing a potentially stronger link to management practices and dispersal pathways by explaining 41% together (Figure 3b). Mobile ARGs were less tied to specific farms (33% vs 43% for all genes) and more responsive to environmental characteristics, indicating greater potential for spread.

Ecological and environmental factors explained a large portion of ARGs trait variance, especially for those potentially in mobile genetic elements, like plasmids, integrons, and transposons (Figure 3c). The spatial proximity was the most influential factor in drug class differences, especially for genes related to resistance to aminoglycosides, tetracyclines, and MLS (random: spatial autocorrelation > 20% variance explained) and indicated that they are more tied to farming practices. Genes conferring resistance to sulfonamides or beta-lactams showed a higher effect of mouse and environmental factors (> 30%) (Figure 3d). Genes corresponding to target replacement and inactivation resistance mechanisms were more influenced by environmental factors (> 35%) (Figure 3e).

To understand the influence of individual factors on the occurrence of different ARGs, we compared the gamma coefficient estimates of the jSDMs (ᵞ) to quantify the effect and directionality of each mouse-related, environmental or climatic factor on ARGs, averaged across their different traits (Figure 4). While most of the factors included in our models had a negative association with the occurrence of ARGs at different agglomeration levels, livestock density and agricultural land were two factors strongly associated overall, although not in all cases with support higher than 75%. We observed a positive association between the surface of agricultural land to genes related to aminoglycoside, MLS, and tetracycline inactivation that could be localised in mobile genetic elements. The presence of ARGs in integrons, as compared to chromosomal ARGs, was strongly and positively associated with higher pig density (ᵞ > 10, support: 0.75). The genes detected in house mice with potential co-localisation in integrons associated with pig density were *sul1*, *aadA24* and *dfrA5*. These genes are linked to conferring resistance against sulfonamide, aminoglycosides and diaminopyrimidine antibiotics, respectively. The presence of ARGs conferring resistance to fluoroquinolone, phenicol and rifamycin (drug class: aromatics) showed a well supported but low decrease with the increase in cattle density (ᵞ < 0, support: 0.95). Clinically and veterinary relevant ARGs conferring resistance to tetracycline were positively associated with high pig density (ᵞ > 0, support: 0.75), as well as other genes related to fusidane and pleuromutilin (drug class: terpenoids; ᵞ > 0, support: 0.75). When we focused on the impact of genes agglomerated by resistance mechanisms, a high density of pigs and cattle was positively associated with genes involved in target replacement mechanisms, including *sul* and *dfr* genes, which could also be mobile (ᵞ > 0, support: 0.75). While pig density was negatively associated with the occurrence of ARGs with target protection mechanisms (ᵞ < -5, support: 0.75), it was positively associated with genes involved in antibiotic inactivation (ᵞ > 5, support: 0.75). The latter included 40 genes for beta-lactamases, such as the AmpC type beta-lactamases *ampC1* and *ampH* (prevalence <10%), as well as the *CbIA-1* gene, which is highly prevalent in mouse resistomes (prevalence >50%). In addition to beta-lactamases, the group of genes involved in antibiotic inactivation influenced by pig density also included the *APH(3’)-IIIa* gene, which is potentially mobile and confers resistance to aminoglycosides (9% prevalence). Particularly, the association between *CblA-1* and pig density was better supported when we analysed the effect on a subset of clinically relevant genes (Supplementary Figure S9).

**Figure 4.**
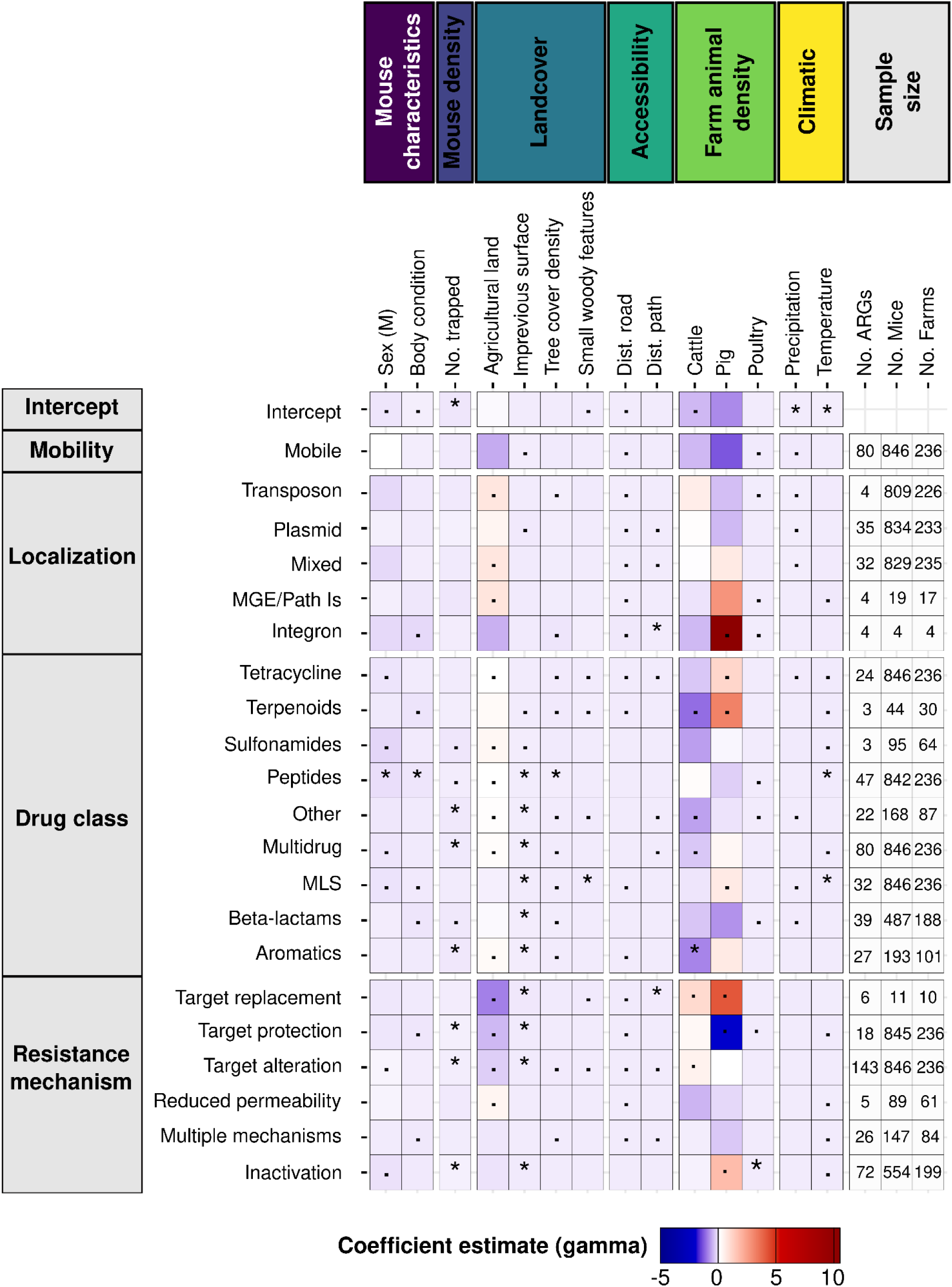
Effect of mouse-related, environmental, farm animals density and climatic variables (columns) on ARG occurrence agglomerated according to their traits (rows). The heatmap shows the coefficient estimate (gamma) for the trait and a given explanatory variable (jSDM). The higher the coefficient estimate, the higher the association of the variable with the ARG trait. The intercept reference level for traits corresponds to ARGs that are non-mobile, located on chromosomes, giving resistance towards the aminoglycoside drug class, and having an efflux resistance mechanism. Symbols in tile indicate the posterior probability of the estimate in the model: ∗ > 0.95 and · > 0.75.

### Effect of environmental characteristics on the ARG compositional structure within house mice microbiomes

We employed the jSDM on the ARG abundance after filtering for rare ARGs to determine the impact of the environmental factors on the compositional structure of ARGs in house mouse microbiomes. The abundance model included 161 ARGs and similarly to the occurrence based jSDM, the diagnostic parameters indicated adequate model convergence (Supplementary Figure S10). The variance in ARG abundance in the mousegut bacterial microbiome was explained on average almost totally by the random effects (88.1%), and relatively few by the fixed effects (11.9%), which is a much higher proportion compared to models predicting ARG occurrence. Similar to the occurrence model, farm spatial proximity (random effect: spatial autocorrelation) was the dominant source of explained variance. This means that spatial proximity of samples led to similar ARG abundance profiles, indicating spatially similar conditions or potential local spread not accounted for in our models. The landcover variables explained about 4%, followed by farm animal density, climatic variables and accessibility, each explaining 2%, and mouse-related characteristics each explained 1%. When focusing only on mobile ARGs, mouse-related, ecological and climatic variables explained about the same amount of variance in mobile genes (13.1%) compared to all genes (11.9%) (Supplementary Figure S11). Considering the little variance in ARG abundance explained these variables, we only detected few supported effects. The temperature showed a mild negative association with ARG abundance (ᵞ < 0, support: 0.95). As in the occurrence-based jSDM, farm animal density did not show a well supported relationship with ARGs in transposons and plasmids (ᵞ > 0, support < 0.75) (Supplementary Figure S12).

### Association of house mouse resistomes with livestock resistomes

Given the association of the occurrence of specific ARGs and gene traits within the house mouse resistome with the density of farm animals, we assessed the similarity between resistomes of livestock to those of house mice in our study. We compared the resistomes of house mice to two previously published datasets: one including manure resistomes from pigs, cattle and chickens in nine European countries, and the other including resistomes of rural and urban pigs and poultry (chickens and turkeys) from Ghana, as a non-european reference. We detected 2,834 ARGs in total considering 269 pig, 238 chicken, 29 cattle and 17 turkey manure metagenomic samples in addition to our 859 house mousegut metagenomes. Overall, we observed that house mouse resistomes were less rich in ARGs than livestock resistomes (Wilcoxon *p*-value adjusted <0.01, effect size *r* >0.2, Supplementary Table S13). To quantify the relative effect of these differences, we fitted a linear model with ARG richness as response. Estimated marginal means confirmed the differences between mice to the other host’s resistomes. However, the difference between the resistomes of mice and pig manure was smaller than the one observed for the other hosts (Contrast_pig-mouse_: 173, SE [6.06], Figure 5a).

**Figure 5.**
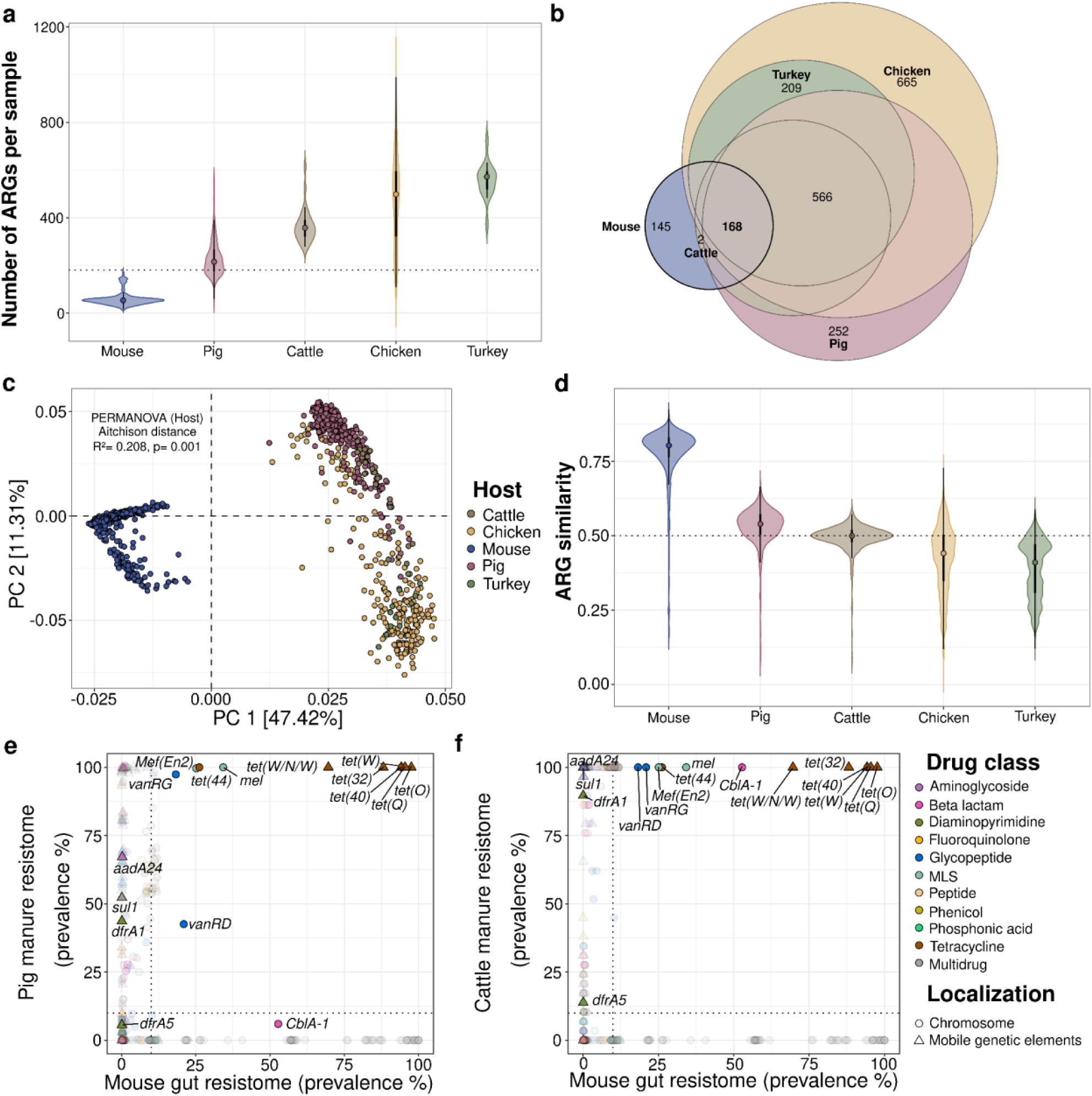
Similarities between the house mouse resistome and livestock manure ARG reservoirs. **(a)** ARG richness in house mice compared to livestock manure. The number of ARGs detected in house mice was significantly lower than in any other host’s manure resistome (Wilcoxon, *p*-value adjusted <0.001). Poultry manure had the highest ARG richness. Pig manure’s resistomes had a smaller difference to mouse resistome richness than for the other hosts (Contrast _pig-mouse_: 173, SE [6.06]). The dashed black line defines the maximum number of ARGs detected in house mice. Each point represents the median per host. Violin represents the distribution of richness per host’s resistome. Solid boxes inside the violin indicate the interquartile range and whiskers extend to minimum and maximum values per host. **(b)** Euler diagram of shared ARGs between house mouse resistomes and different livestock manure resistomes; numbers next to the host labels indicate the number of unique genes for that particular host. A total of 168 genes were shared between the mouse resistome and the other hosts. The livestock resistomes shared 566 ARGs among them. **(c)** Principal component analysis (PCA) showing the dissimilarity in resistome composition among hosts. Dissimilarity is based on Aitchison distances between samples on centered log ratio transformed ARG abundance. Each dot represents an individual resistome. **(d)** Comparison of ARG composition similarity between mouse resistomes and each of the other host’s resistomes. ARG similarity was calculated as 1 - Aitchitson distance between two samples. Similarity values of 1 represent identical ARG profiles, while those equal to zero indicate completely different resistomes. The similarity values between each mouse resistome and each host are presented. Pig and cattle manure’s resistomes had a higher compositional similarity to mouse resistomes than to the poultry hosts (contrast _pig-mouse_: -0.264, SE [0.0098], contrast _cattle-mouse_: -0.310, SE [0.013]). The dashed black line indicates a similarity value of 0.5. Points represent the median resistome similarity between mouse and a given host. Violin represents the distribution of similarity to mice resistomes per host. Solid boxes inside the violin indicate the interquartile range and whiskers extend to minimum and maximum similarity values. **(e)** Prevalence of ARGs in the house mouse gut and pig manure metagenomes. While the gene *CbIA-1* had higher prevalence in mouse guts than in pig manure indicative of house mouse promotion, three genes that can be localised in integrons (*aadA24*, *sul1* and *dfrA1*) were more prevalent in pig manure than in mice, indicating a promotion of those genes in pig manure. **(f)** Prevalence of ARGs in the house mouse gut and cattle manure metagenomes. The gene *CbIA-1* prevalent in more that 50% of the mousegut metagenomes could be detected in all the cattle manure resistomes, similar to those genes that can be localised in integrons. For both environments, six *Tet* genes had a prevalence above 75% indicating co-promotion in such environments. In panels e and f, labels are shown for ARGs with prevalence higher than 25% in either of the hosts and those genes that could be localised in integrons. The shape indicates whether the gene can be localised in the chromosome (circles) or in mobile genetic elements (triangles) and thus suggest mobility potential. Dashed lines indicate a cutoff at 10% prevalence to highlight genes associated with one of the hosts or in both.

To determine the potential connectivity between house mouse resistomes and livestock manure, we calculate the number of ARGs shared between mice and each livestock host, as well as the composition similarity. Considering the total number of different ARGs detected (N= 2,834) in the distinct hosts, the resistomes of mice shared a small fraction of genes (n= 168) with those of livestock manure and only had a significant overlap with cattle’s manure resistome (Fisher’s exact test, n= 174 shared genes, *p*-value adjusted <0.01, Figure 5b). The house mouse resistome showed a distinctive ARG composition compared to the resistomes of livestock manure (PERMANOVA, R^2^= 0.21, *p*-value <0.01, Figure 5c). Resistome similarity was higher between resistomes of mice and pig and cattle manure than between resistomes from poultry hosts (contrast _pig-mouse_: -0.264, SE [0.0098], contrast _cattle-mouse_: -0.310, SE [0.013], Figure 5d).

We compared the gene prevalence from both pigs and cattle against the prevalence in house mice resistomes to determine genes that could be promoted by a specific host or in both host’s resistomes. Those genes present in ≥10% of both mouse and livestock host metagenomes were classified as co-promoted, while genes with prevalence ≥10% in only one of the hosts were classified as host-specific-promoted. When mouse and pig manure prevalences were compared, we observed that 11 genes were co-promoted, particularly tetracycline resistance genes (*Tet*) associated with mobile genetic elements. In contrast, three genes encoded in integrons (*aadA24*, *sul1* and *dfrA1*) were promoted in pig manure, but still detected in mice resistomes (Figure 5e). The gene *CbIA-1* encoding a non-mobile beta-lactamase has higher prevalence in mice gut than in pig manure indicative of house mouse promotion. A similar picture arose from the comparison between mouse and cattle manure, with *tet* and integron associated genes being co-promoted, but with the difference that *CblA-1* is also co-promoted in both cattle manure and mice, suggesting the circulation of such genes between manure and house mice (Figure 5f).

## Discussion

Our study showed that the occurrence of antimicrobial resistance genes in the microbiomes of natural populations of generalist species, such as house mice, is strongly influenced by land use characteristics. Here, we provided a novel framework to assess how environmental gradients influence ARG composition and the potential interactions that facilitate gene mobility among microbial communities, by conceptualizing the ARGs as metacommunities within host-associated microbiomes sampled across transects within heterogeneous landscapes. We demonstrated that environmental variables, particularly livestock farming intensity, explained a substantial proportion of the variation in ARG occurrence, especially among genes carried by mobile genetic elements. Notably, high pig farming density was closely associated with integron-encoded sulfonamide resistance genes, as well as with genes conferring resistance to tetracyclines and beta-lactams. These findings demonstrate that specific ARGs in the resistomes of wild house mice are also found in livestock manure, highlighting the exchange of resistance genes between agricultural and natural ecosystems.

Our metagenomic surveillance revealed that the resistomes of house mice are primarily composed of genes of limited clinical or veterinary concern, largely involved in multidrug resistance mechanisms that are ubiquitous in bacterial genomes of both host-associated and environmental origin^56,57^. This pattern is consistent with our previous 16S-based predictions of chromosomally encoded ARGs in house mice^58^ specifically, and with wild rodent resistomes in a broader sense^59^. However, we were able to detect genes potentially co-localised with mobile genetic elements. A previous study in wild house mice from urban environments reported a wide distribution of ARGs conferring resistance to macrolide–lincosamide–streptogramin (MLS) antibiotics, fluoroquinolones, tetracyclines, and beta-lactams^37^. While our house mouse resistomes included some of these previously reported genes and others of high priority^50^, they were detected here at low abundance and prevalence compared to other environments like the gut and livestock microbiome or wastewater systems. One explanation for the low prevalence could be the reduced abundance or absence of their typical bacterial hosts. Families of commensal gut bacteria such as *Enterobacteriaceae*, *Enterococcaceae*, and *Staphylococcaceae*, which are usually involved in mediating ARG transmission within host-associated microbiomes^22,60,61^, were poorly represented in our house mouse samples. In other wild rodent species, culture-based analyses have revealed elevated levels of resistance among *Enterobacteriaceae*^35^. In contrast, wild cotton mice (*Peromyscus gossypinus*) inhabiting polluted environments with microbiomes dominated by *Desulfovibrionaceae*, *Pseudomonadaceae*, and *Helicobacteraceae* showed high abundance and prevalence of multidrug resistance genes, with only a few genes associated with last resort antibiotics^62^. Conversely, wild rodents from rural areas in Germany appeared to play only a minor role in the transmission of resistant *Escherichia coli* to the environment^63^, consistent with our broader characterization of the house mouse resistome. Together, these observations suggest that the resistomes of wild house mice, similar to those of other rodents, are dominated by environmentally ubiquitous ARGs. The house mouse ARG composition was largely shaped by the bacterial taxa present in their microbiomes, and both bacterial and ARG composition could reflect the degree of anthropogenic exposure and potential for gene exchange across ecological boundaries.

Beyond microbial interactions within the host that may drive the selection of specific ARGs, we focused on identifying ecological drivers of the house mouse resistome. Given that both biotic and abiotic factors shape microbial communities through natural selection, and selective pressures may vary from habitat to habitat^64^, we hypothesized that rodents’ gut microbial communities and associated genetic repertoires are shaped by exposure to diverse environmental conditions. We observed that spatial localization of each farm exerted a strong influence on the overall ARG composition. Ecological and environmental variables also explained a substantial proportion of the variability in ARG occurrence, exceeding the effects of host density and sex, consistent with previous observations in the human gut resistome^19^. These patterns likely reflect anthropogenic factors that promote antibiotic resistance in bacteria within the mouse gut. Environmental and climatic factors were influential for genes potentially co-localised on mobile genetic elements and therefore more likely to be transferred through horizontal gene transfer. This suggests that mobile genes may be promoted by changes in agricultural intensity or land management practices unlike chromosomal, non-mobile ARGs, which are less susceptible to external environmental pressures. In external environments such as soil and water, conventional agricultural management practices, including the use of manure as fertiliser, herbicides or compounds containing heavy metals, increase the prevalence of ARGs and mobility between bacteria by horizontal gene transfer^65–68^. We found that extensive agricultural land use was positively associated with potentially mobile genes, suggesting that those genes may be selected either through the direct impact of stress-inducing xenobiotics that reach wildlife via water and soil, or present in their microbiome through the invasion of bacteria carrying ARGs selected in the agricultural environment, as previously observed in humans^69,70^. Moreover, It has been shown that cross-species ARG transmission between livestock and wild mammals is mediated by their co-occurrence with mobile genetic elements^71^. The dissemination of ARGs between different environments, and between host-associated and environmental microbiomes increases with the intensity of anthropogenic activities^72^. Thus, the influence of anthropogenic agricultural and farming practices are persistent on the house mouse resistome.

The use of antibiotics for food production has long been recognized as a strong contributor to antimicrobial resistance in the human microbiome, independently of clinical antibiotic consumption^73–75^. By modelling the effect of agricultural surface and livestock density on the mouse resistome, we aimed to test whether the effect of antibiotic use for food production can be quantifiable not only in humans but also in other species inhabiting human-impacted environments. Similar to agricultural practices, livestock density, particularly of pigs, was strongly associated with potentially mobile ARGs within house mouse microbes. The association between livestock density and mouse resistome can reflect two interconnected processes: dissemination of mobile ARGs through fecal pollution and epidemiological connectivity through host movement and interactions^76^. As livestock resistomes are considered “hotspots” of resistance genes, the dissemination of ARGs through manure occurs at a larger extent than from human, sewage and soil sources^13,77^. Pig manure is highly enriched in mobile ARGs and represents a relevant source for spreading ARGs to the environment^78–81^. The movement of mice between habitats and social interactions likely increase their exposure to sites contaminated with livestock faeces or with other rodents carrying ARGs^34,82^. Consistent with our results, other studies detected that wild rodents, and even larger species such as wild boar (*Sus scrofa*) from rural areas with high livestock density in Germany, carry *E. coli* resistant to antibiotics commonly administered in pigs, poultry and cattle suggesting livestock to wildlife transmission^39,63^. Overall, interactions between agricultural management practices, livestock density and mouse mobility may have significant impacts on the composition and mobility of resistomes in wild house mouse populations.

We observed a strong association of pig density to highly prevalent genes conferring resistance to tetracycline (*tet*), which could be traced in both the house mouse resistome and in pig and cattle resistomes from Germany and other European countries. Tetracyclines have been historically used in livestock husbandry in large-scale intensive farming to prevent diseases and promote animal growth^83,84^. In Germany, tetracyclines remain among the most widely administered antibiotics, together with beta-lactams^85,86^. Despite its regulation intended to minimise selection of resistant bacteria^87^, residues of tetracyclines and their bacterial transformation products persist in livestock manure and fertilisers at concentrations that still represent an ecological risk and have antibacterial activity^88,89^. Thus, *tet* genes are persistent and highly abundant in environments with fecal or manure pollution. These genes are considered part of the cross-environment core resistome and useful for the monitoring of the total resistome diversity in the environment^15,90,91^. In our study, we detected six *tet* genes that may reflect the impact of livestock in the house mouse resistome, as they were enriched in both environments. The transmission of these genes from livestock to wildlife likely results from their co-localisation on mobile genetic elements, which facilitates efficient spread through horizontal gene transfer across bacteria^92,93^. Tetracycline ARGs are known to be promiscuous and their capacity to transfer between phylogenetically distant bacterial hosts^14^. Our results indicate that house mouse resistomes are strongly impacted by livestock density, and that mobile *tet* genes represent good markers for tracking ARG transmission between farm environments and wildlife species.

Genes encoding for commensal beta-lactamases, like *CblA-1* and *sul1* were additional markers of anthropogenic and livestock influence identified in the house mouse resistome. The *sul1* gene, although not specifically promoted in house mouse resistomes, provides interesting insights into potential ways of resistance transmission. *sul1* is a mobile and highly prevalent gene in different environments, particularly human-impacted surface waters and wastewater commonly used for agricultural irrigation^94,95^. In riverine environments, animal feeding operations have been identified as a relevant source of *sul1*^96^, highlighting how this gene may serve as a marker of connections between farming activities, agricultural practices, and the dissemination to broader environments, including wildlife hosts such as house mice. We also detected that the C*blA-1* gene was highly promoted in house mouse resistomes. Although *CblA-1* is not a mobile gene and is restricted to the genus *Bacteroides*^97,98^, this *CblA-1* cephalosporinase is highly prevalent in the human gut microbiomes from Western countries and may exacerbate antimicrobial resistance infections if transferred to pathogens^99^. The presence of this gene likely reflects human or cattle fecal pollution, further supporting its role as an indicator of anthropogenic impact in wildlife resistomes. Taken together, genes as *sul1* and *CblA-1* can be complementary indicators of livestock-derived and anthropogenic pollution within the house mouse resistome.

Our findings reveal that antimicrobial resistance in wildlife is not an isolated phenomenon but a measurable reflection of anthropogenic activity across agricultural landscapes. We integrated metagenomic screening and ecological data to demonstrate the impact of the external environment on the resistome of natural populations of house mice, allowing the identification of specific genes that trace this impact. The transmission dynamics between livestock, environment and other mammal hosts, such as house mice, can be explained by the influx of selective agents and bacteria carrying ARGs from the farming environment into surrounding habitats, where ARGs could be maintained within wildlife microbiomes. Unlike other environments such as wastewater, polluted soil and the gut of humans and livestock, the house mouse resistome is characterised by several regulators that do not themselves confer a resistant phenotype and fewer of the high-priority genes identified by the WHO. In this respect, mice are not the most relevant carriers and are definitely not the most practical samples for surveillance purposes. However, our analysis clearly shows that their resistomes reflect anthropogenic impact, particularly with regard to genes carried by mobile genetic elements. Therefore, we identified mobility as an important trait for assessing how farm animals and agricultural practices influence the resistance profiles of natural house mice populations in surrounding environments. This study broadens our understanding of the transmission of antimicrobial resistance in human-impacted environments and supports the development of strategies to monitor the mobility and dissemination of ARGs and resistant bacteria into the environment and wildlife species.

## Acknowledgments

This work was supported by the DFG Research Infrastructure NGS_CC (project 407495230) as part of the Next Generation Sequencing Competence Network (project 423957469). NGS sequencing was carried out at the Competence Centre for Genomic Analysis (Kiel). VHJD was supported by DFG FO1279/6-1, MG, AP and SCMF were supported by DFG KR4266/4-1 and HE7320/5-1. VHJD and SKFS are supported through the JPIAMR-EMBARK (F01KI1909A) and JPIAMR-SEARCHER (01KI2404B) projects funded by the Bundesministerium für Bildung und Forschung (BMBF).

We thank Moritz Wenzler-Meya for his support in preparing the spatial layers. We thank the JPIAMR-EMBARK and SEARCHER consortia for the feedback on the results that improved this manuscript.

## Author contribution

**MG:** methodology, investigation, formal analysis, writing - review editing; **AP**: methodology, investigation, data curation, formal analysis, writing - review editing; **EH:** conceptualisation, funding, supervision, resources, writing - review editing; **SKFS:** conceptualisation, funding, resources, writing - review editing; **SKS:** conceptualisation, funding, supervision, project administration, writing - review editing (equal); **SCMF:** methodology, investigation, writing - review editing; **VHJD:** conceptualisation, methodology, investigation, data curation, formal analysis, writing - review editing, funding, supervision

## Data availability

All raw sequence data are deposited and available online at the European Nucleotide Archive with an accession number PRJEB102564. Additional data for the analysis and the code used for visualization and statistical analysis are available at https://github.com/mgicquel/mus_musculus_de_amr (non-static version under development) and will be archived at the EcoDyn github account and Zenodo upon publication.

