## Supplementary for "Anthropogenic land use exerts selection pressures on the resistome of a wild rodent"

### Supplementary material

**Supplementary Table S1.** Classification of antimicrobial resistance genes (ARGs) into drug classes and broader drug class groups for the association analysis in the joint species distribution models (jSDMs). To address the sparse representation of certain drug classes, ARGs were grouped into broader drug class groups based on chemical structure and mechanism of action. This table details the reclassification of individual drug classes to their corresponding broader groups, along with the number of ARGs identified per class and per group.

| Drug class group | Drug class | Number of ARGs per class | Number of ARGs per group |
|---|---|---|---|
| Aminoglycosides | aminoglycoside | 59 | 59 |
| Aromatics | fluoroquinolone | 23 | 29 |
| | phenicol | 3 | |
| | rifamycin | 3 | |
| Beta-lactams | beta-lactam | 8 | 39 |
| | carbapenem | 4 | |
| | cephalosporin | 27 | |
| MLS | elfamycin | 7 | 32 |
| | lincosamide | 6 | |
| | macrolide | 18 | |

| | | | |
|---|---|---|---|
| | streptogramin | 1 | |
| Multidrug | multidrug | 81 | 81 |
| Other | aminocoumarin | 4 | 22 |
| | diaminopyrimidine | 4 | |
| | mupirocin | 1 | |
| | nitroimidazole | 1 | |
| | nucleoside | 1 | |
| | phosphonic acid | 10 | |
| | zoliflodacin | 1 | |
| Peptides | glycopeptide | 25 | 48 |
| | peptide | 23 | |
| Sulfonamides | sulfonamide | 3 | 3 |
| Terpenoids | fusidane | 1 | 3 |
| | pleuromutilin | 2 | |
| Tetracyclines | tetracycline | 24 | 24 |

**Supplementary Table S2.** Clinically relevant groups of antimicrobial resistance genes (ARGs).

| Gene group | Description | Resistance mechanism | Drug class |
|---|---|---|---|
| CblA | CblA beta-lactamase | antibiotic inactivation | cephalosporin |
| CepA | CepA beta-lactamase | antibiotic inactivation | cephalosporin |
| CfxA | CfxA beta-lactamase | antibiotic inactivation | cephamycin |
| CTX-M | CTX-M beta-lactamase | antibiotic inactivation | cephalosporin |
| KPC | KPC beta-lactamase | antibiotic inactivation | monobactam, carbapenem, cephalosporin, penam |
| SHV | SHV beta-lactamase | antibiotic inactivation | carbapenem, cephalosporin, penam |
| TEM | TEM beta-lactamase | antibiotic inactivation | penam, monobactam, cephalosporin, penem |
| CfiA | CfiA beta-lactamase | antibiotic inactivation | carbapenem |

| | | | |
|---|---|---|---|
| IMP | IMP beta-lactamase | antibiotic inactivation | carbapenem, cephalosporin, cephamycin, penam |
| NDM | NDM beta-lactamase | antibiotic inactivation | carbapenem,cephalosporin,cephamycin,penam |
| VIM | VIM beta-lactamase | antibiotic inactivation | carbapenem, cephalosporin, cephamycin, penam |
| ACT | ACT beta-lactamase | antibiotic inactivation | carbapenem, cephalosporin, cephamycin, penam |
| ampC | ampC-type beta-lactamase | antibiotic inactivation | cephalosporin, penam |
| CMY | CMY beta-lactamase | antibiotic inactivation | cephamycin |
| OXA | OXA beta-lactamase | antibiotic inactivation | carbapenem, cephalosporin, penam |
| AAC(3) | AAC(3) | antibiotic inactivation | aminoglycoside |
| AAC(6') | AAC(6') | antibiotic inactivation | aminoglycoside |
| ANT(3") | ANT(3") | antibiotic inactivation | aminoglycoside |
| ANT(4') | ANT(4') | antibiotic inactivation | aminoglycoside |
| ANT(6) | ANT(6) | antibiotic inactivation | aminoglycoside |
| APH(2") | APH(2") | antibiotic inactivation | aminoglycoside |
| APH(3') | APH(3') | antibiotic inactivation | aminoglycoside |
| APH(3") | APH(3") | antibiotic inactivation | aminoglycoside |
| APH(6) | APH(6) | antibiotic inactivation | aminoglycoside |
| vanA | vanA/vanB | antibiotic target alteration | glycopeptide |
| qnr | Quinolone resistance protein | antibiotic target protection | fluoroquinolone |
| MCR | Phosphoethanolamine transferase | antibiotic target alteration | peptide |
| pmr | Phosphoethanolamine transferase | antibiotic target alteration | peptide |
| Fos | Fosfomycin thiol transferase | antibiotic inactivation | phosphonic acid |
| cat | Chloramphenicol acetyltransferase | antibiotic inactivation | phenicol |

| | | | |
|---|---|---|---|
| tet | tetracycline-resistant ribosomal protection protein | antibiotic target protection | tetracycline |
| tet | tetracycline inactivation enzyme | antibiotic inactivation | tetracycline |
| Cfr | Cfr 23S ribosomal RNA methyltransferase | antibiotic target alteration | oxazolidinone, streptogramin, phenicol, lincosamide |
| Erm | Erm 23S ribosomal RNA methyltransferase | antibiotic target alteration | macrolide, lincosamide, streptogramin |
| lnu | Lincosamide nucleotidyltransferase | antibiotic inactivation | lincosamide |
| Ere | Macrolide esterase | antibiotic inactivation | macrolide |
| mph | Macrolide phosphotransferase | antibiotic inactivation | macrolide |
| arr | rifampin ADP-ribosyltransferase | antibiotic inactivation | rifamycin |
| sul | Sulfonamide resistant | antibiotic target replacement | sulfonamide |

**Supplementary Table S3.** Environmental layers used for modelling ARG occurrence in the mouse gut microbiome. All spatial layers contain continuous data. Year refers to the year of origin of the data. For distance to roads and paths, the references provided refer to the original layers of respective streets and paths used to calculate the distances. Corine Land Cover layer was used to calculate the proportion of agricultural fields (*agri*).

| Variable | Environmental layer | Units | Year | Original resolution |
|---|---|---|---|---|
| agri | CORINE land cover[1] | proportion of agricultural land (land use class 2) | 2018 | vector data |
| imperv | Imperviousness[1] | proportion of impervious surface per raster cell | 2018 | 10m raster |
| tcd | Tree cover density[1] | proportion of area occupied by trees per raster cell | 2018 | 10m raster |
| swf | Small woody features[1] | proportion of area occupied by linear structures such as hedgerows, as well as patches of woody features per raster cell | 2018 | 100m raster |
| dist_roads, dist_paths | Distance to roads, distance to paths [2] | streets and paths linear, rasterised and transformed into distances in meters | 2018 | vector data |
| catdens, pigdens, poudens | Livestock[3] | Livestock units of cattle, pigs and poultry, transformed into density per hectare | 2020 | vector data |

| | | | | |
|---|---|---|---|---|
| prec | Precipitation[4] | Monthly total precipitation (l/m2 or mm) | 2015-2022 | 1km raster |
| temp | Temperature[4] | Monthly average air temperature (1/10 °C) | 2015-2022 | 1km raster |

[1] https://land.copernicus.eu/en/products/ (Downloaded 06.02.2024)

[2] obtained from GeoBasis-DE / BKG (2022): https://gdz.bkg.bund.de/index.php/ (Downloaded 07.02.2024)

[3] obtained from the Thünen Atlas: Landwirtschaftliche Nutzung Winterweizen / Landwirtschaftliche genutzte Fläche 2020, https://atlas.thuenen.de/webspace/agraratlas/agraratlas/index.html?LP=2 (Downloaded 18.09.2023)

[4] obtained from the Deutscher Wetterdients: https://opendata.dwd.de (Downloaded 05.11.2024)

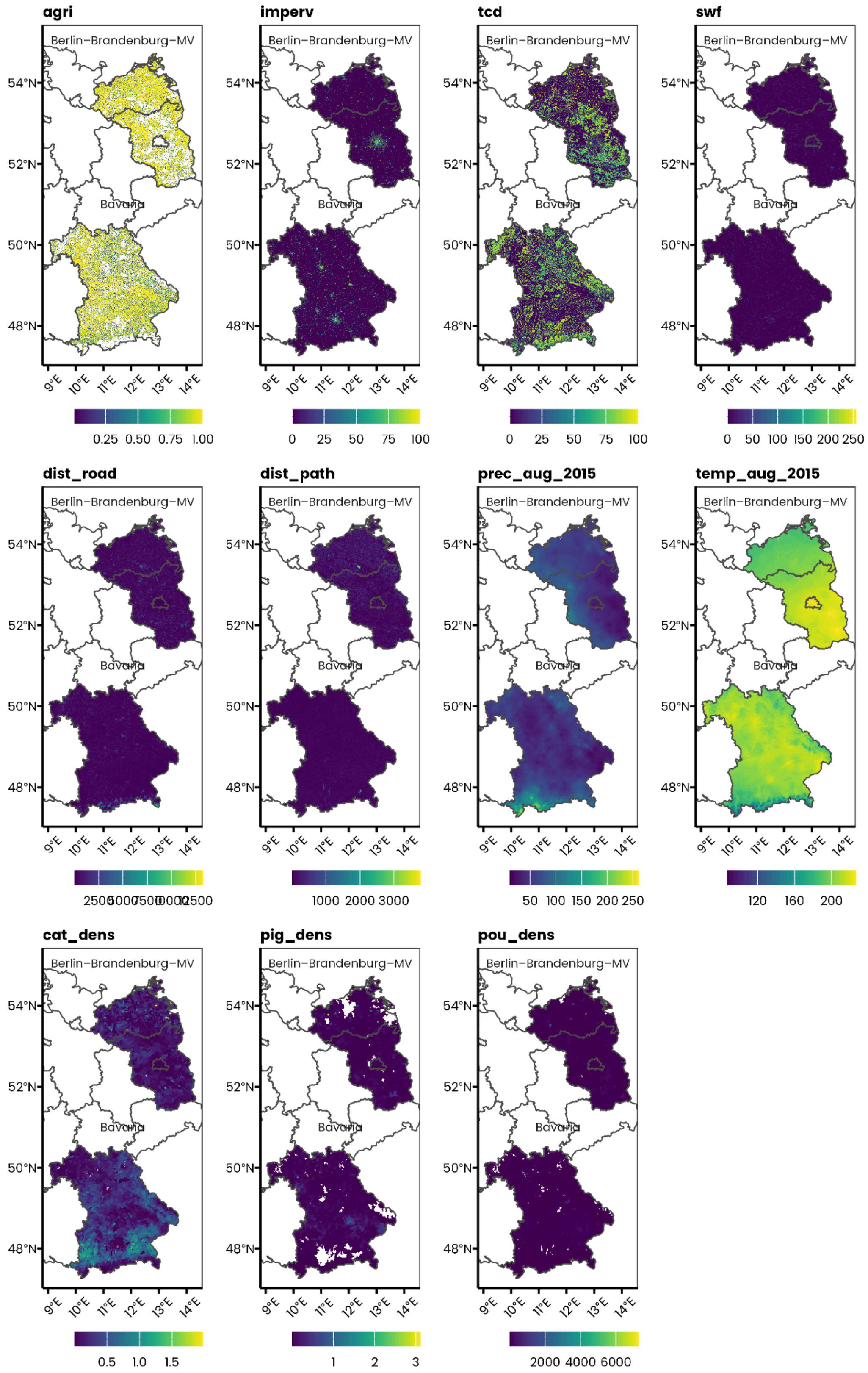


**Supplementary Figure S4.** Maps of environmental layers in the sampled regions (Berlin-Brandenburg-Mecklenburg-Vorpommern and Bavaria). For precipitation and temperature, we only displayed the values recorded during the year 2015.

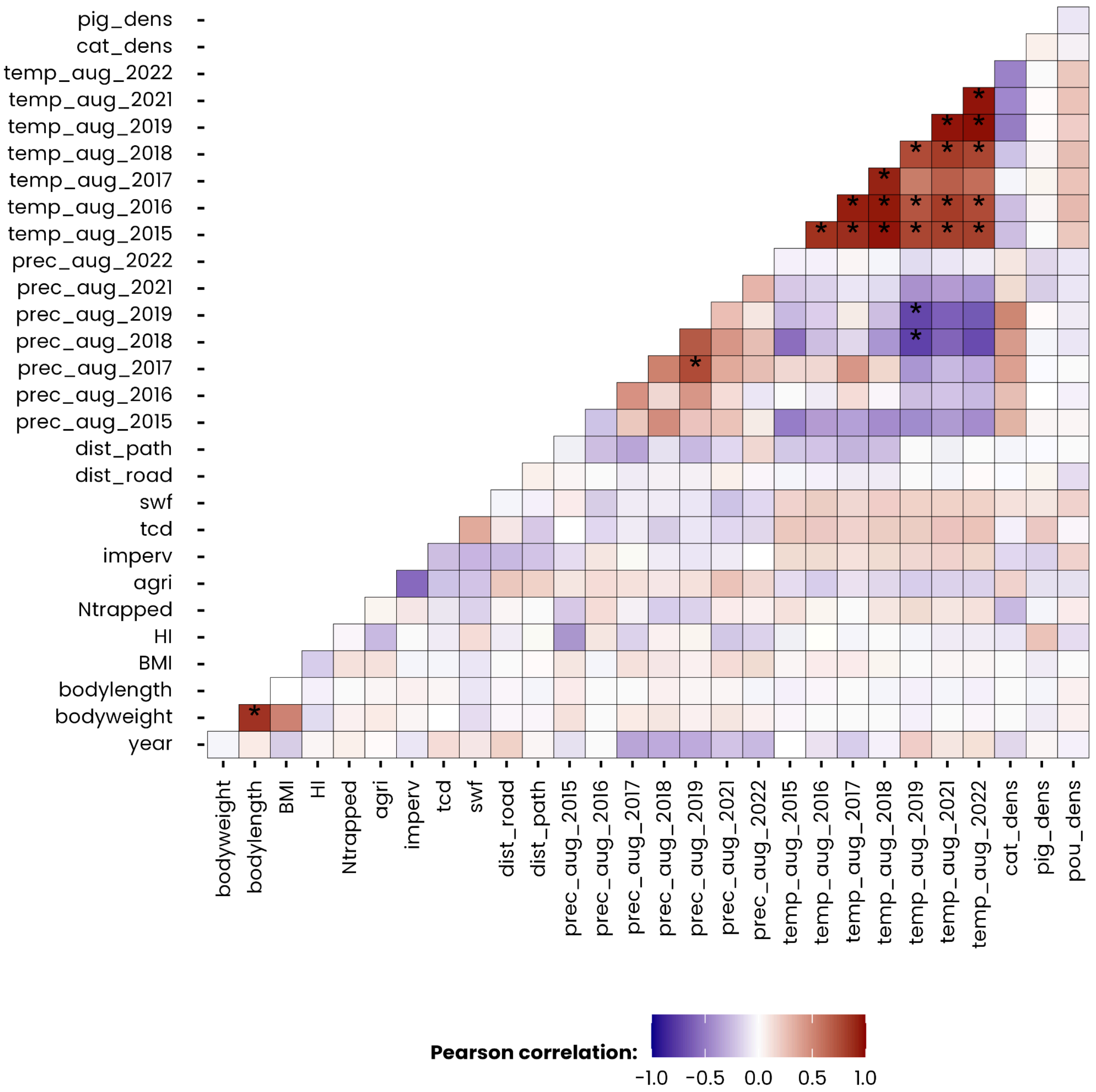


**Supplementary Figure S5.** Pearson correlation matrix showing the pairwise linear relationships between the explanatory variables. Colours indicate the strength and direction of the correlation, with blue representing negative correlations and red representing positive correlations. Symbol ∗ in tile indicates correlation values |r| > 0.7.

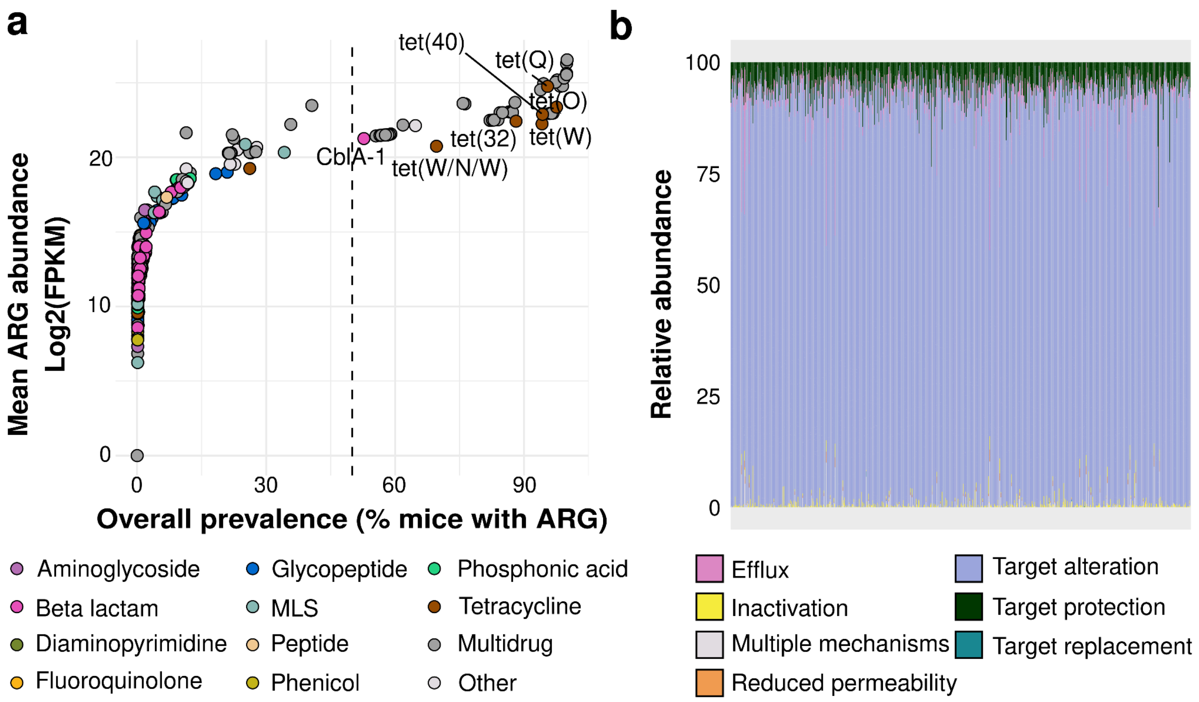


**Supplementary Figure S6.** Overall prevalence and abundance per ARG. **(a)** Mean abundance and prevalence of ARGs in the colon content metagenomes from house mice. In total, 55 genes had an overall prevalence of more than 50% and a mean abundance over 2,500,000 FPKM across the metagenomes. Only six of these highly prevalent genes confer resistance to two relevant drug classes: tetracycline and beta-lactam. **(b)** Individual mouse relative abundances by resistance mechanism. Genes related to target alteration and protection were consistently more abundant in all mouse metagenomes.

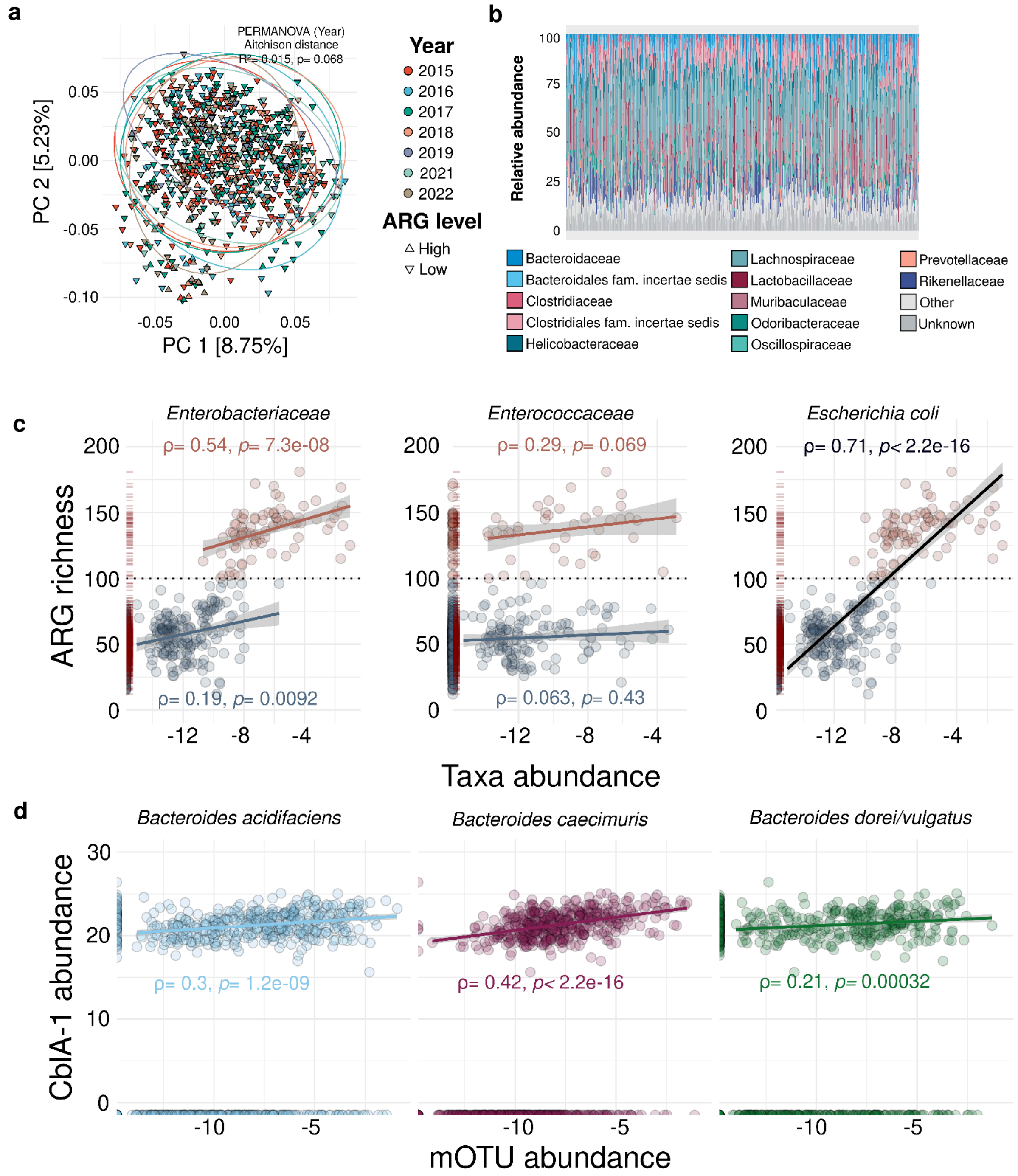


**Supplementary Figure S7.** General overview of the colon microbiomes from house mice and their relationship to ARG levels. **(a)** Principal component analysis (PCA) showing dissimilarity in microbial composition between years of sampling. Each point represents the colon microbiome of a sample. Distances between points reflect biological gene composition dissimilarity based on Aitchison distances. Points and ellipses are coloured by year of sampling. Shape represents whether an individual metagenome has higher or lower than 100 ARGs predicted. **(b)** Individual mouse relative abundances of operational taxonomic (mOTUs) units agglomerated by bacterial families. mOTUs from the Lachnospiraceae family were consistently more abundant and prevalent in all mouse metagenomes (Mean relative abundance= 25.8% [95%CI: 24.8-26.8], Prevalence= 99.8%). **(c)** Spearman's rank correlations between microorganisms commonly associated with resistance and ARG richness per mouse. Commensal gut families (*Enterobacteriaceae*, *Enterococcaceae*) explain differences in ARG levels in mice. Higher abundance of such families is related to higher levels of resistance. *Escherichia coli* has a strong correlation to the total ARG richness. **(d)** The abundance of the *CblA-1* gene coding for a Class A beta lactamase is related to *Bacteroides* sp. Within the house mouse microbiome. *B. caecimuris* abundance has a stronger correlation compared to the other two *Bacteroides* species with prevalence above 50%. Taxa abundance is expressed as the log2 of the relative abundance, and for the gene *CblA-1* the abundance is expressed as log2 FPKM.

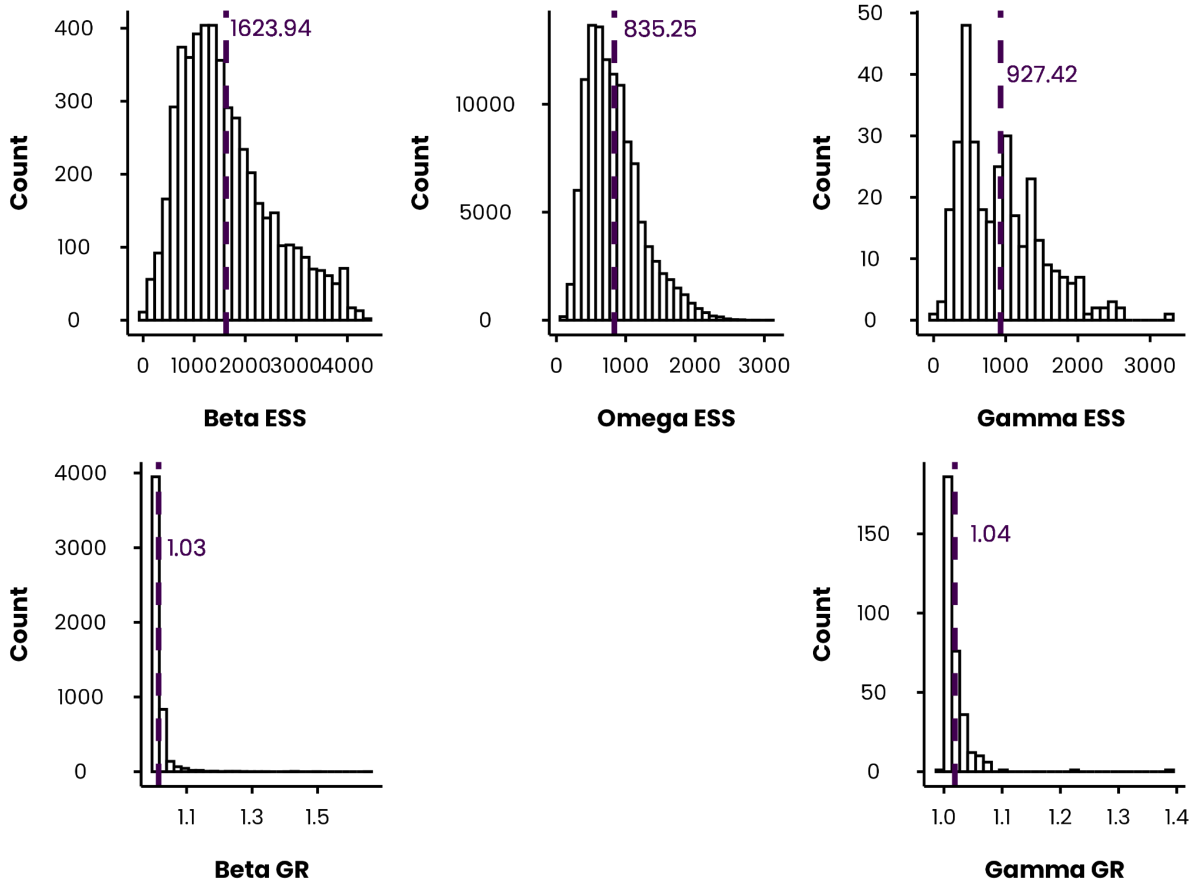


**Supplementary Figure S8.** Convergence diagnostics for the ARG presence-absence joint species distribution model (jSDM). The top row shows the distribution of effective sample sizes (ESS) for monitored parameters, with the dashed purple line indicating the average ESS. The bottom row shows the distribution of Gelman–Rubin (GR) statistics, with the dashed purple line marking the average GR value. Model convergence: The average effective sample sizes for all parameters are high (>800), indicating that the model estimates robust parameters and exhibits low autocorrelation and good mixing. Gelman-Rubin diagnostic values are acceptable for betas (1.03) and gammas (1.04), which means chains converge well. It could not be compiled for omegas for structural reasons (huge matrices for each sample). Model fit metrics: Pseudo-$R^2$ (Tjur $R^2$) = 0.13 , AUC = 0.91, $R^2$ sp = 0.17, $R^2$ site = 0.72

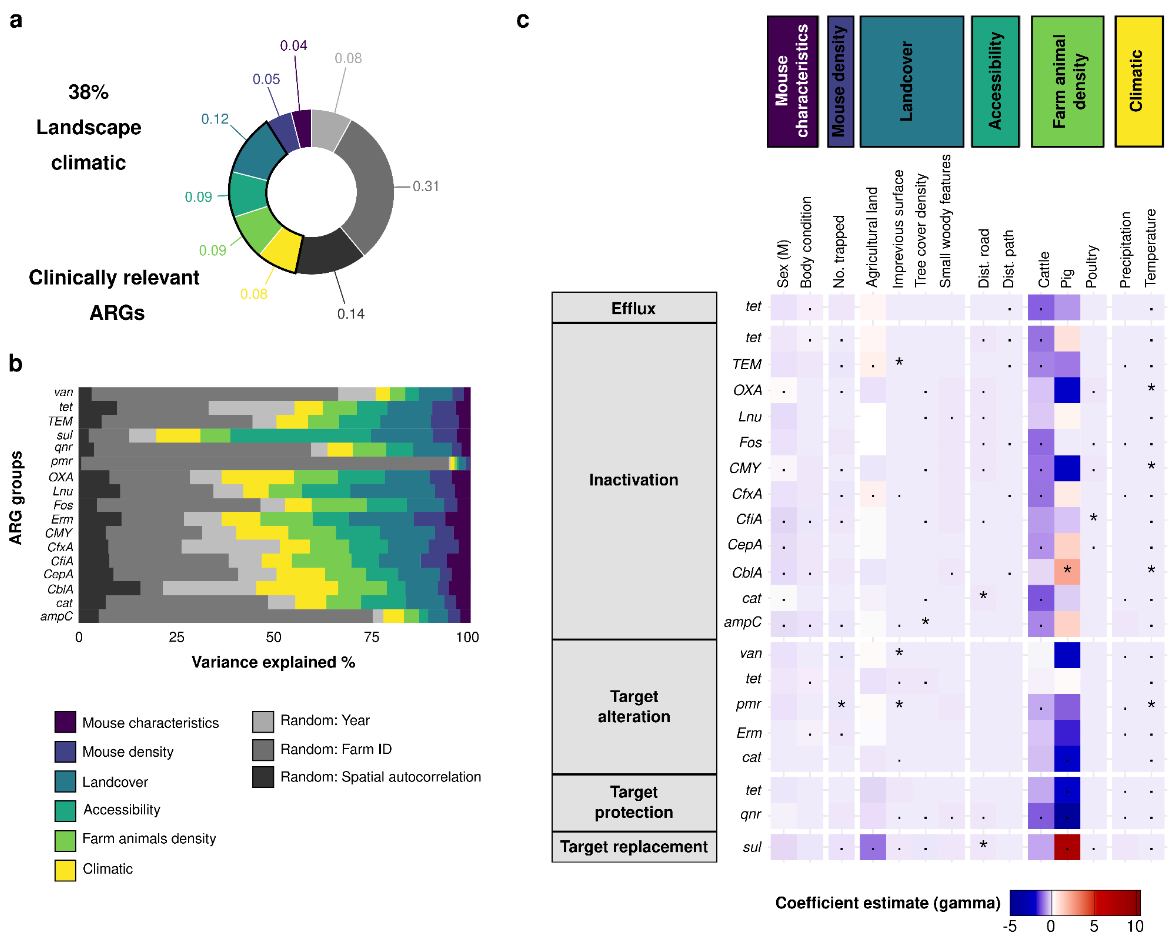


**Supplementary Figure S9.** Variance partitioning showing the relative contribution of mouse characteristics, environmental factors, and random effects, on the explained variation in clinically relevant ARG occurrence in mouse gut microbiomes. **(a)** Landcover accounts for the higher proportion of variance on these genes **(b)** Variance partitioning in occurrence averaged for each ARG group. For *sul* genes landscape characteristics are particularly relevant and account for more than 70% of occurrence variance. **(c)** The heatmap shows the coefficient estimate (gamma) for each ARG group and a given explanatory variable of the jSDM. Cattle and pig density are explanatory variables with stronger associations compared to other variables in the model. *sul* genes with target replacement mechanism and the *cblA-1* beta-lactamase are strongly and positively associated with pig density ($\gamma > 5$, support: 0.75). *tet* genes are distinctively associated with pig density, while those related to antibiotic inactivation have a positive association, those involved in efflux and target protection mechanisms are negatively associated, and especially the latter with a strong effect ($\gamma < 0$, support: 0.75). The higher the coefficient estimate, the higher the association of the variable with the ARG group. Symbol in tile indicate the posterior probability of the estimate in the model: $\ast > 0.95$ and $\cdot > 0.75$.

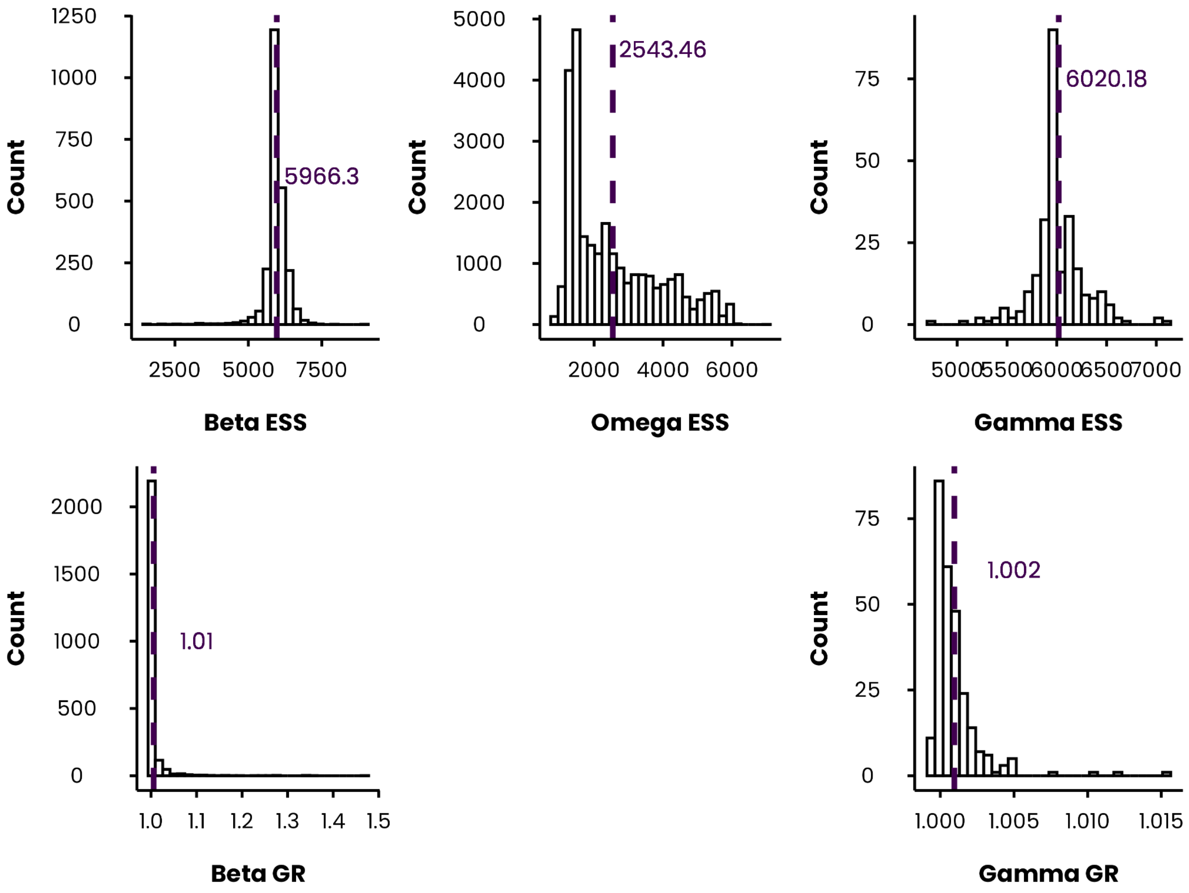


**Supplementary Figure S10.** Convergence diagnostics for the jSDM ARG abundance model. The top row shows the distribution of effective sample sizes (ESS) for monitored parameters, with the dashed purple line indicating the average ESS. The bottom row shows the distribution of Gelman–Rubin (GR) statistics, with the dashed purple line marking the average GR value. Model convergence: The average effective sample sizes for all parameters are high (>2500), indicating that the model estimates robust parameters and exhibits low autocorrelation and good mixing. Gelman-Rubin diagnostic values are acceptable for betas (1.01) and gammas (1.002), which means chains converge well. Model fit metrics: Pseudo-$R^2$ (Tjur $R^2$) = 0.22 , AUC = 0.796, $R^2$ sp = 0.22, $R^2$ site = 0.81

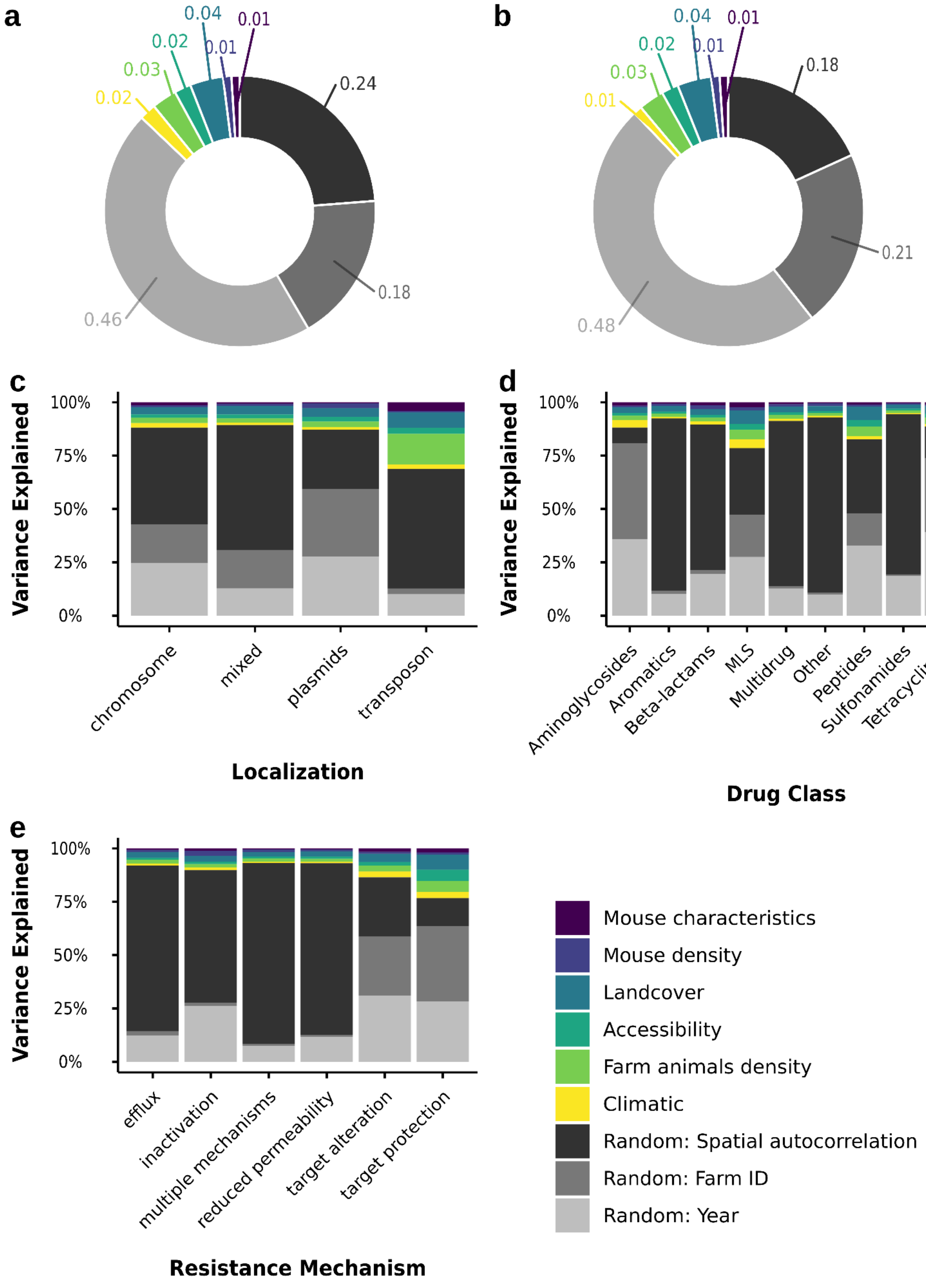


**Supplementary Figure S11.** Variance partitioning showing the relative contribution of mouse characteristics, environmental factors, and random effects, on the explained variation in ARGs abundance in mouse gut microbiomes **a)** for all genes and **b)** only for mobile genes. **(c-e)** Variance partitioning in abundance averaged for each ARG trait (localization, drug class and resistance mechanism).

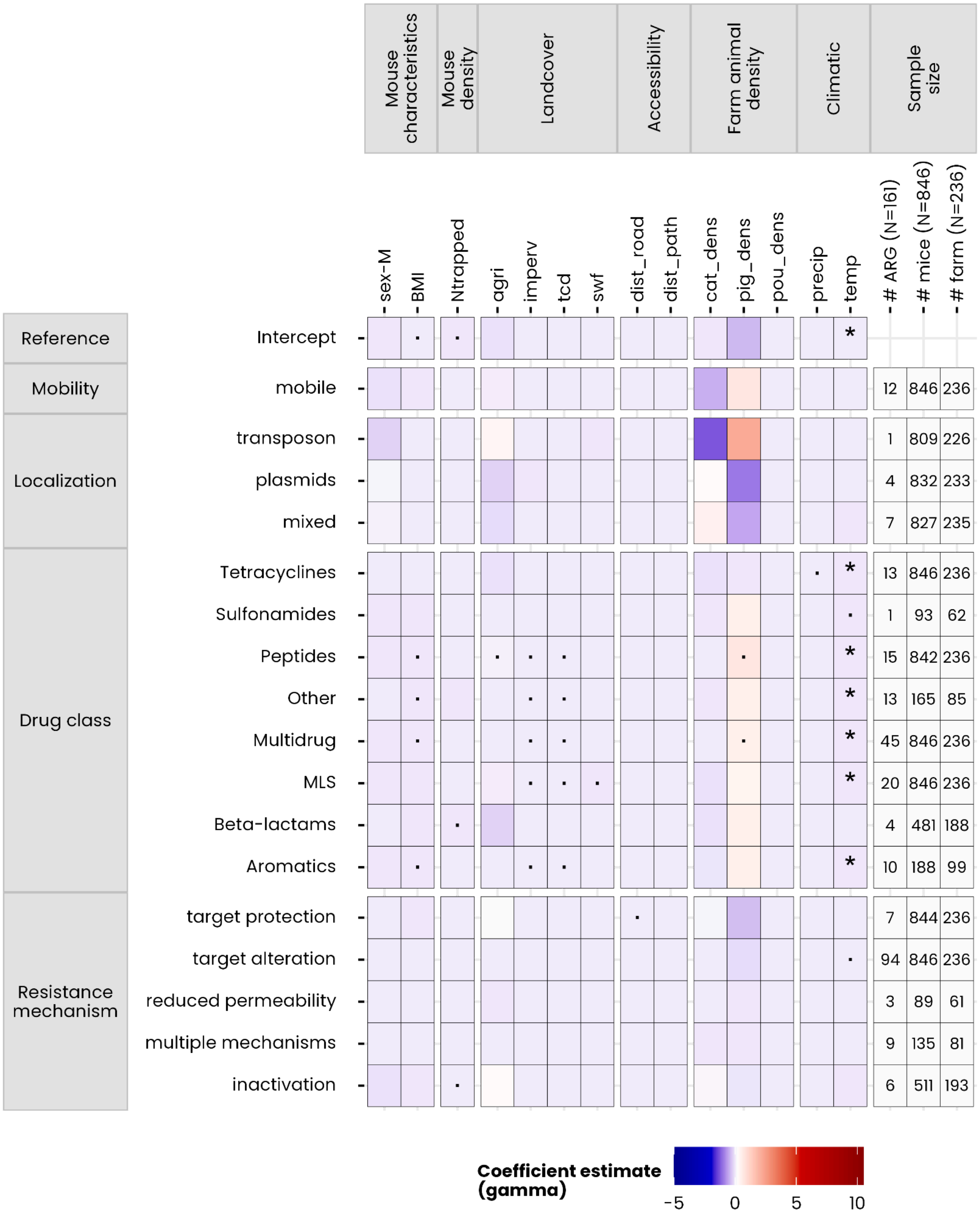


**Supplementary Figure S12.** Effect of mouse-related, environmental, farm animal density and climatic variables on ARG abundance agglomerated according to their traits. The intercept reference level for traits corresponds to ARGs that are non-mobile, located on chromosomes, giving resistance towards the aminoglycoside drug class, and having an efflux resistance mechanism. Symbol in tile indicate the posterior probability of the estimate in the model: ∗ > 0.95 and · > 0.75.

**Supplementary Table S13. Differences in ARG richness between house mice and livestock manure**

| Host 1 | Host 2 | P-value adj. | Significance | Effect size | Contrast |
|---|---|---|---|---|---|
| House mice (n=859) | Cattle (n=29) | 4.73E-19 | **** | 0.31 | 306±16.4 |
| | Chicken (n=238) | 1.29E-121 | **** | 0.71 | 416±6.4 |
| | Pig (n=269) | 6.10E-132 | **** | 0.73 | 173±6.1 |
| | Turkey (n=17) | 1.56E-12 | **** | 0.24 | 497±21.2 |
